# Horizontal gene transfer of the functional archaellum machinery to Bacteria

**DOI:** 10.1101/2025.02.02.636118

**Authors:** Shamphavi Sivabalasarma, Najwa Taib, Clara L. Mollat, Marie Joest, Stefan Steimle, Simonetta Gribaldo, Sonja-Verena Albers

## Abstract

Motility in Archaea is driven by a nanomachinery called the archaellum. So far, archaella have been exclusively described for the archaeal domain; however, a recent study reported the presence of archaellum gene clusters in bacterial strains of the SAR202 clade (Chloroflexota). Here, we show that *bona fide* archaellum gene clusters are widespread in several members of the Chloroflexota, which in turn lack any bacterial flagellar components. Analysis of archaellum encoding loci and predicted structures show remarkable similarity to the archaellum machinery. Moreover, using cryoEM single particle analysis, we solved the structure of the bacterial archaellum from *Litorilinea aerophila*, demonstrating the successful expression and assembly of this machinery in Bacteria and its function in swimming motility. Finally, a phylogenomic analysis revealed two horizontal gene transfer events from euryarchaeal members to Chloroflexota. In summary, our study demonstrates that a functional and assembled archaellum machinery can be successfully exchanged between the two prokaryotic domains.

## Introduction

Throughout the three domains of life, organisms evolved different macromolecular machineries for cell motility and propulsion^1^. In Archaea motility is driven by the archaellum^2^. Despite the functional resemblance to the bacterial flagellum, the archaellum belongs to the type IV filament superfamily (TFF). This superfamily comprises diverse macromolecular machineries with different functions, with the type IV pilus being the namesake of this superfamily (Figure 1a). In Archaea, the TFF family diversified into different subtypes, each comprising a functionalized cell surface structure such as the ups pilus, the aap pilus, or the bindosome^3–6^. The hallmark of the TFF superfamily is a four-protein core decorated with machinery-specific accessory proteins, with the archaellum being the only rotary member^3,7^. The archaellum filament is composed of the archaellin ArlB (or paralogue ArlA) in single or multiple copies processed by a class III signal peptidase^8,9^. ArlI is the ATPase that powers the assembly and rotation of the archaellum filament, likely in interaction with the membrane platform protein ArlJ^10–12^ (Figure 1a). ArlH is a KaiC homolog and can modulate its oligomeric state and interaction with ArlI through autophosphorylation^13,14^. ArlF and ArlG form the stator complexes necessary for non-futile rotation and torque generation^15,16^ (Figure 1a). In Thermoproteota, ArlX forms a ring-like complex that likely supports the core machinery of ArlJIH and the stator complex^17^. Euryarchaeota instead have ArlCDE, which are essential to transfer the signals from the chemotaxis system to the archaellum motor^4,17,18^. All these genes are organized within a cluster of 7-11 genes that are all essential for motility and archaellation^19,20^. The difference in operon organization and the presence of ArlCDE distinguishes the clusters in *arl1* and *arl2* (formerly *fla1* and *fla2*) between Thermoproteota and Euryarchaeota^21^.

**Figure 1:**
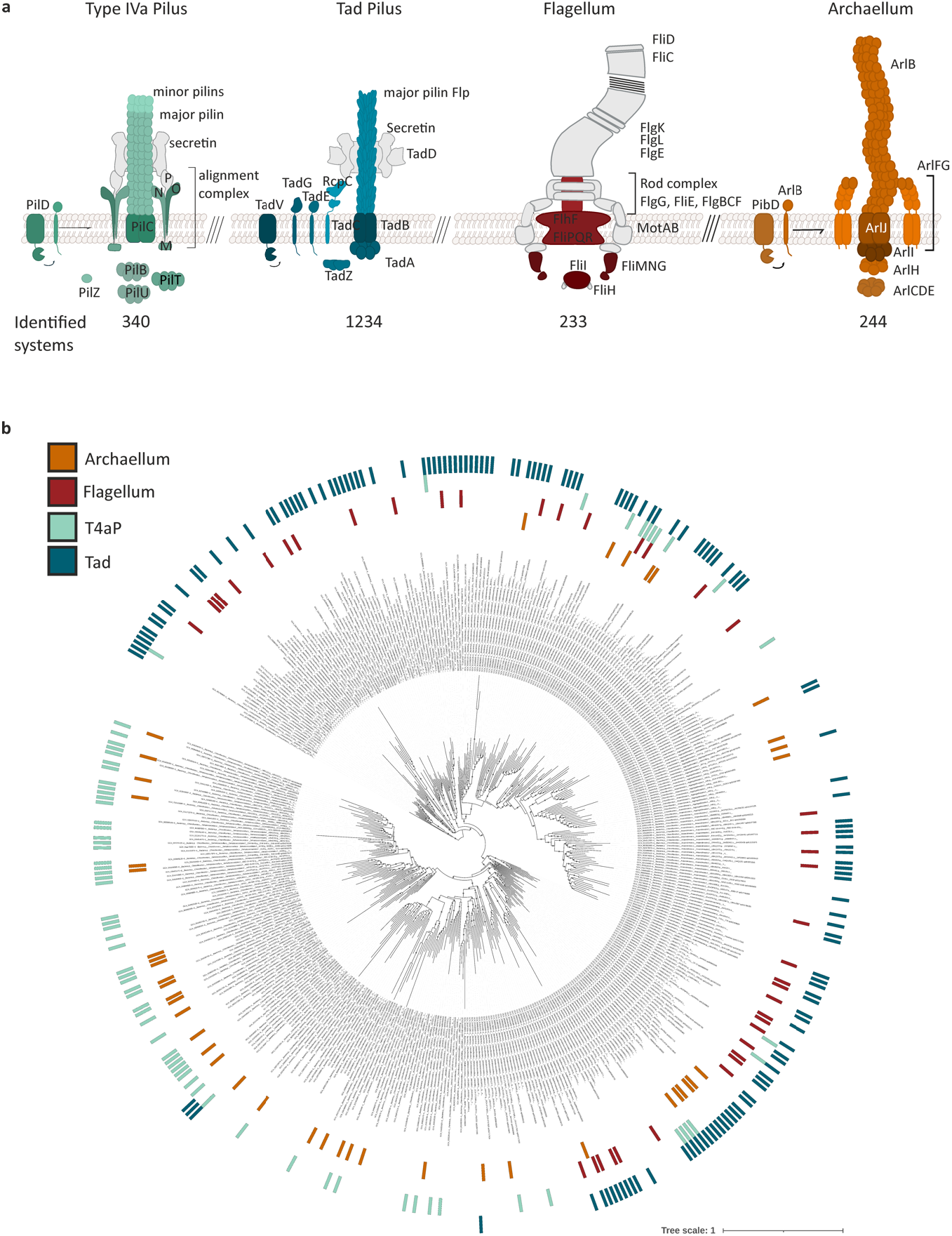
Cell surface machineries in Chloroflexota. **a** Schematic representation of searched macromolecular systems and associated proteins. Each system with the corresponding proteins was searched in Chloroflexota genomes using MacSyFinder2. The present subunits are colored accordingly, and gray subunits were not found. Of the 3780 genomes searched, 233 potentially flagellated members were found. 1234 encoded for a Tad pilus, and 339 have loci for a Type IVa pilus. 244 genomes encoded for a complete archaellum machinery. **b** Phylogenetic tree of all Chloroflexota genomes analyzed with the present TFF and flagellum systems. Archaellum machineries are found in most order of Chloroflexota. Tree scale as indicated.

So far, it was thought that the archaellum was solely employed by Archaea for swimming motility, while Bacteria use the flagellum^1^. In a recent study, the first members of the SAR202 clade, marine Bacteria belonging to Chloroflexota, were cultivated^22^. Analysis of their genome sequence showed that these Bacteria encode for a *bona fide* archaellum machinery with several tandem copies of the filament encoding gene *arlB*^22^. However, isolated SAR202 clade strains lack cell surface filaments indicative of an assembled archaellum^22^. In addition, metagenomic sequences of Chloroflexota from aquifer sediments have been reported to encode an archaellum machinery missing the crucial prepilin/archaellin peptidase PibD, suggesting it likely does not assemble into a functional archaellum^23^.

In this study, we screened a locally maintained database of curated prokaryotic genomes for the presence of secretion systems, TFF machineries and flagellum-related genes using MacSyFinder2^24^. While most bacteria do not harbor any archaellum-related genes, several Chloroflexota encode a *bona fide* archaellum machinery. An in-depth bioinformatic analysis of 3780 genomes of Chloroflexota revealed the presence of the archaellum operon in 244 members, including the cultivated species *Litorilinea aerophila*. We demonstrated the expression and assembly of a functional archaeal-like archaellum structure by *L. aerophile*, which is used for swimming motility. CryoEM single-particle analysis of the purified archaella filaments showed remarkable structural similarity to the archaellum. The preservation of the structural features of the bacterial archaellum, analogous to that of its archaeal equivalent, suggests a conserved mechanism for archaellum-driven swimming in *L. aerophila*. Phylogenomic analysis of the archaellum cluster in archaea and bacteria revealed two horizontal gene transfer events from euryarchaeal members of Methanotecta to Dehalococcoides. The monoderm envelope architecture found in Chloroflexota might have eased the successful incorporation of the archaellum machinery into the bacterial envelope.

## Results

### Taxonomic distribution of the archaellum machinery in Archaea and Bacteria

Previous studies on Chloroflexota metagenomes and the first cultivated member of the SAR202 clade (Ca. *Lucifigimonas marina*) indicated the presence of genes related to archaella in Bacteria^22,23^. A locally maintained database of archaeal and bacterial genomes with a representative taxon sampling was screened for the presence of TFF and flagellar macromolecular systems using MacSyFinder2^24^. We found that Bacteria lack the genes coding for the archaellum machinery, with the notable exception of a few members of Chloroflexota. We, therefore, focused on a more detailed analysis of 3780 available genomes of Chloroflexota. We found that 244 genomes appear to code for the archaellum machinery, while 233 genomes contain flagellar-related genes. Around 1234 members encode for a Tad pilus, and we detected 340 genetic loci coding for a type IVa pilus (Figure1ab). When the archaellum machinery genes were present, flagellar-related genes were not found; however, some genomes additionally code for a Tad or type IVa pilus (Figure 1b).

When present, the flagella machinery in these Chloroflexota seems to lack components compared to other flagellated Bacteria. In fact, a comparison with the genome of *Tepidiforma thermophila*, the only member of the Chloroflexota for which flagella have been shown to be assembled, revealed that the genes for stator complexes and flagellar filaments are lacking and likely being missed due to the employed MacSyFinder model^58^. Since most genomes are not fully assembled and flagellar components are rarely found within a single genetic locus, detecting the remaining flagellar components with certainty was challenging. The Tad pilus was found in roughly a third of the genomes analyzed, revealing its widespread in Chloroflexota. The identified Tad and type IVa pili systems lacked secretins, corroborating the absence of an outer membrane in this phylum^59^ (Figure 1ab, Supplementary table 3). Remarkably, archaellum loci are the most complete having all genes necessary for the expression of a fully functional machinery (Figure 1a, Supplementary Table 3). When the genes for the core machinery (*arlJIH*) and *arlB* were identified, the accessory genes for the stator complex (*arlFG*) and the archaeal switch complex (*arlCDE*) were generally also present. In conclusion, this indicates the presence of many bona fide archaellum machineries within Chloroflexota.

### Analysis of archaellum encoding loci in Chloroflexota

We identified around 244 archaellum clusters in Chloroflexota across different orders. Selected archaellum clusters from members of the SAR202 clade, Anaerolineales, Dehalococcoidales, and Thermofilales were compared with archaellum clusters from known archaellated and motile archaeal species^19,60–65^. All archaellum-encoding Chloroflexota possess the filament subunit *arlB* in single or multiple copies (Figure 2a, Supplementary Table 3). Downstream of the *arlB*(s), we identified genes encoding the archaellum machinery within one genetic locus. Like the euryarchaeal archaellum systems, most Chloroflexota encode *arlCDE* homologs instead of the Thermoproteota-specific *arlX*. However, in contrast to most Euryarchaea such as *Haloferax volcanii* or *Pyrococcus furiosus*, Chloroflexota harbor the *arlG and arlF* genes in reverse order (Figure 2). This dimorphism in gene organization is characterized as the *arl1* and *arl2* locus organization. While most Euryarchaea have an *arl1* organization with the order *arlFG*, Thermoproteota and some Euryarchaea have *arl2* with the *arlGF* configuration^21^. In contrast to Archaea, some Chloroflexota have a duplication of the stator gene *arlF* and the order of the *arlHIJ* genes is seemingly more variable than in Archaea, nevertheless the genes were always present within the same genetic locus. The class III signal peptidase, necessary for the processing of the filament protein ArlB, called PibD in Archaea and PilD in Bacteria, was identified in these Chloroflexota, but outside of the archaellum cluster locus (Figure 2, Supplementary table 3). Additional genes were found near the archaellum machinery, and *Litorilinea aerophila* specifically encodes a *pilN* and a *pilO* homolog with a LysM domain (Figure 2). In summary, the analysis of the archaellum gene loci in Chloroflexota revealed that the arrangement of machinery genes is comparable to that in Archaea and contain all essential genes for archaellation, suggesting that these surface structures might be functionally assembled in Bacteria.

**Figure 2:**
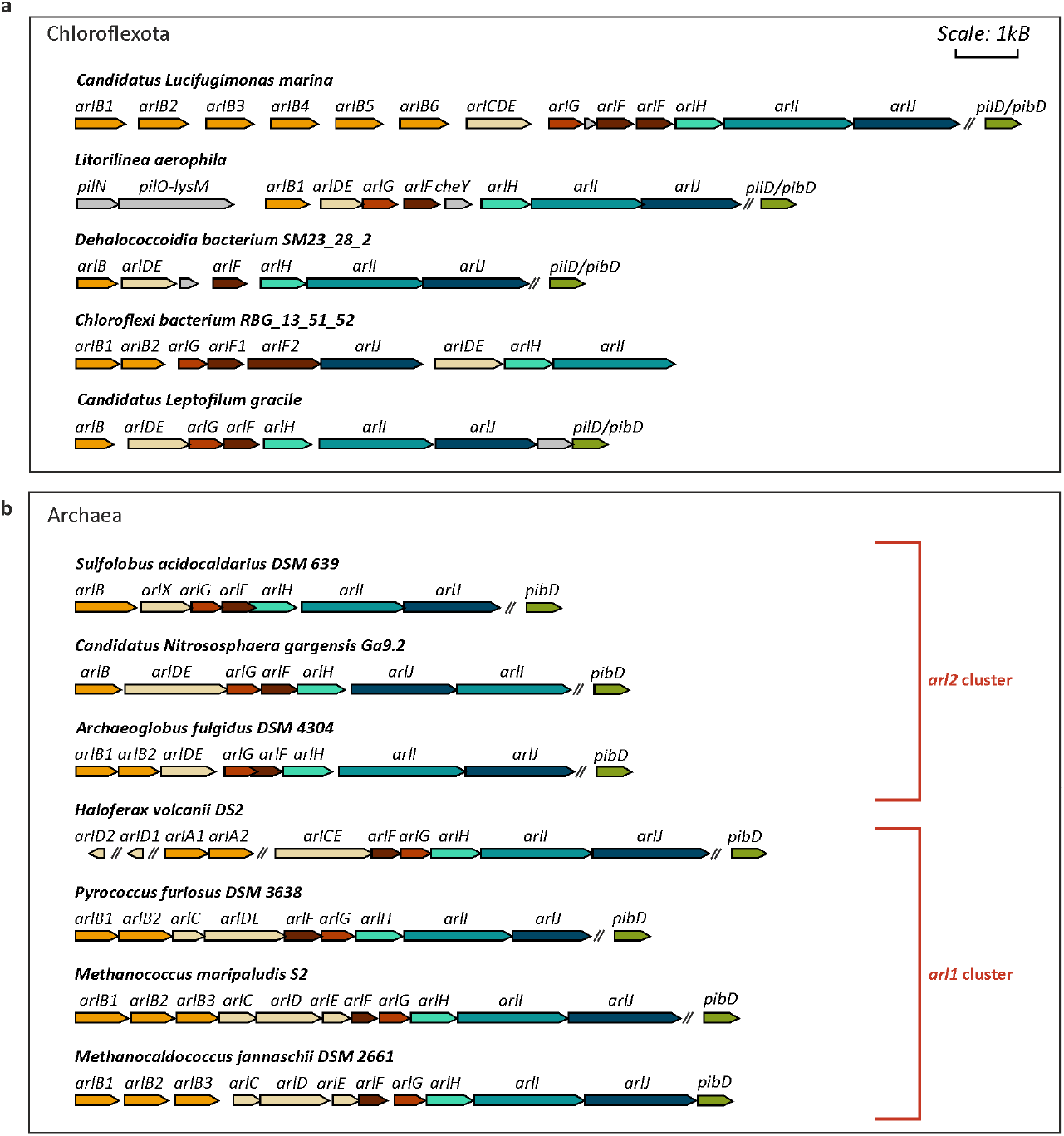
Schematic representation of identified archaellum loci in Chloroflexota in comparison to archaeal archaellum loci. Exemplary identified archaellum loci in Chloroflexota (**a**) and Archaea (**b**). Chloroflexota of different orders harbor archaellum machinery genes. The machinery is complete and has all the necessary genes for archaellation and motility. The order of the genes and their homology is more similar to the *arl2* cluster found in *S. acidocaldarius, Ca. N. gargensis* and the euryarchaeon *A. fulgidus*. Homologous genes are indicated in the same color.

### Structure-guided bioinformatic analysis of the core machinery suggests an archaellum-like rotary mechanism

ATP hydrolysis of ArlI through its interaction with ArlJ drives the archaellum filament assembly and rotation^10^. To assess whether the bacterial archaellum rotates similarly to its archaeal counterpart, we compared the predicted structures of the core machinery proteins

ArlJ and ArlI from *Litorilinea aerophila*. ArlI, the ATPase necessary for assembly and rotation, has a fold and domain organization similar to that of crenarchaeal ArlI from *S. acidocaldarius* (Figure 3)^10^. The C-terminal domain that harbors the ATP binding and hydrolysis site is highly conserved (Figure 3b). The N-terminal domain showed higher variability, but the structure and organization of the domain resembled archaeal ArlI (Figure 3ab). The presence of the archaellum ATPase-specific N-terminal three-helix bundle was striking and suggested its dual function for assembly and rotation as described for the archaeal motor ATPases^10^. Comparison to euryarchaeal ArlI of *M. jannaschii* showed that the bacterial ArlI has a more extended N-terminus with an additional 36 amino acids that are specific to Euryarchaea (Figure 3a). ArlJ encodes the polytopic membrane platform protein with approximately 7-9 transmembrane helices. Few experimental information is available on this protein; however, the predicted structure of bacterial ArlJ shows high similarity to the crenarchaeal ArlJ (Figure 3a). Like archaeal ArlJ, bacterial ArlJ has conserved positively charged residues at the cytosolic site, which are likely crucial for interaction with the ATPase ArlI (Figure 3a)^12^. This striking similarity to the archaeal ArlJI hints at a similar mode-of action for both proteins in acting as core rotary machinery for the archaellum^11^.

**Figure 3:**
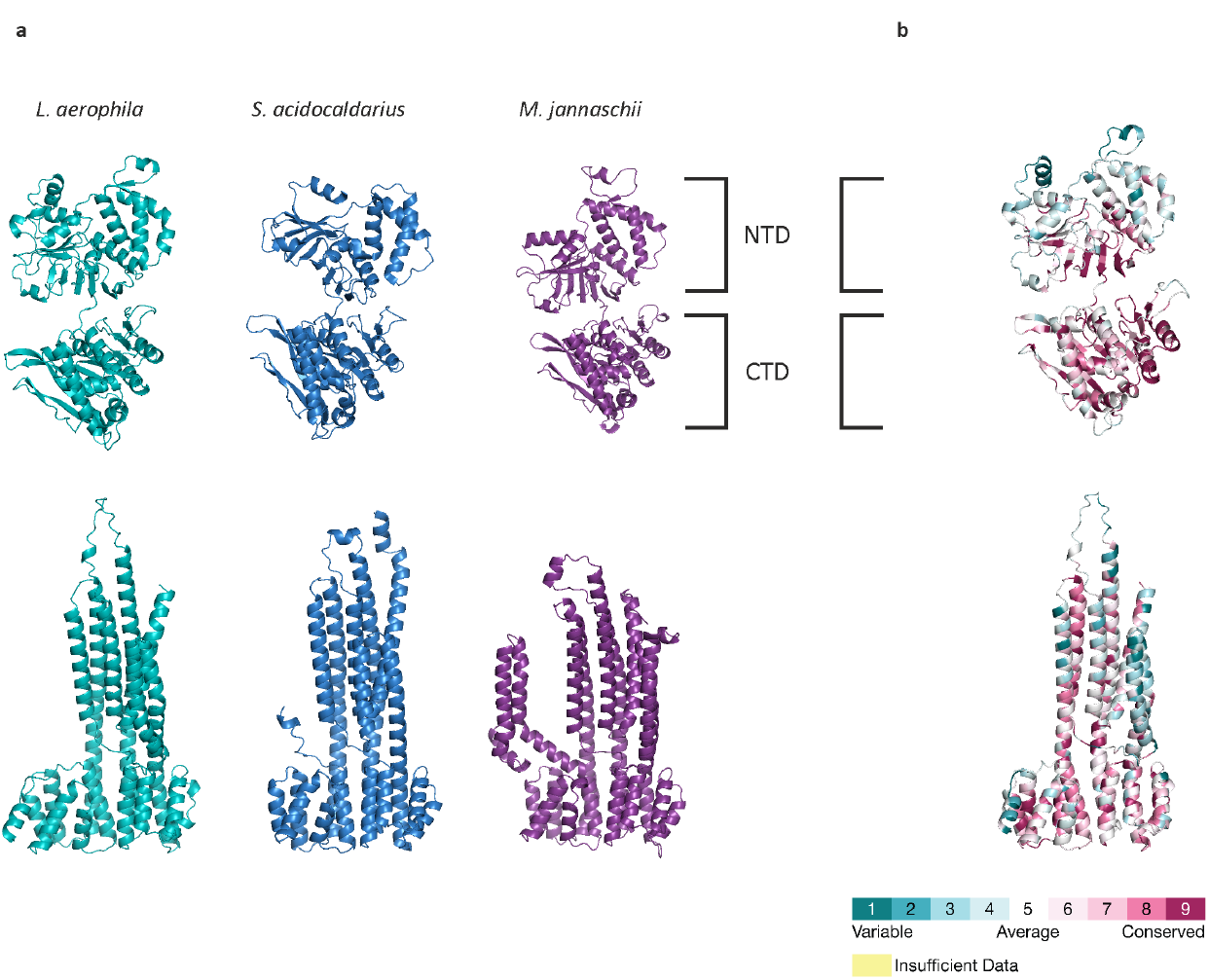
Structure-guided analysis of the core archaellum machinery. **a** Alphafold3 predicted structures of the motor protein ArlI and ArlJ of the bacterial organism *L. aerophila*, the euryarchaeon *M. jannaschii* and *S. acidocaldarius* (pdb: 4ii7 for ArlI). A comparison of all three structures reveals remarkable structural similarity and similar domains in the N-terminal domain (NTD) and C-terminal domain (CTD). **b** Conservation of residues in ArlI and ArlJ of bacterial archaella machineries was calculated using ConSurf. Scale bar indicates calculated conservation scores per residue.

### Litorilinea aerophile, an archaellated bacterium

Our bioinformatic analyses, along with structural predictions of proteins related to the archaellum, suggest that Bacteria of the phylum Chloroflexota might assemble a functional archaellum. To verify this hypothesis, we cultivated the thermophilic filamentous bacterium *Litorilinea aerophila* (Caldlineales)^66^. Although being described as nonmotile, the presence of a complete archaellum cluster suggests the ability of this bacterium to assemble a functional archaellum.

The *L. aerophila* archaellum locus encodes all components necessary for archaellation and a *cheY* homolog (Figure 2a). Additionally, *pilO* and *pilN* homologues are encoded in the genomic neighborhood of the machinery genes (Figure 2, Supplementary Figure 10). The cells were grown in marine broth medium for five days as previously described^67^. After five days the cells grew into long multicellular filaments with individual cell segments. Furthermore, shorter filaments of cells were observed similar to those described for *C. islandicus*^68^ (Figure 4). Electron microscopic analysis of the cells revealed segmentation of the filamentous multicellular bacterium in individual cells (Figure 4). Cell poles were analyzed for archaella filaments, but none were found (Figure 4 upper panel).

**Figure 4:**
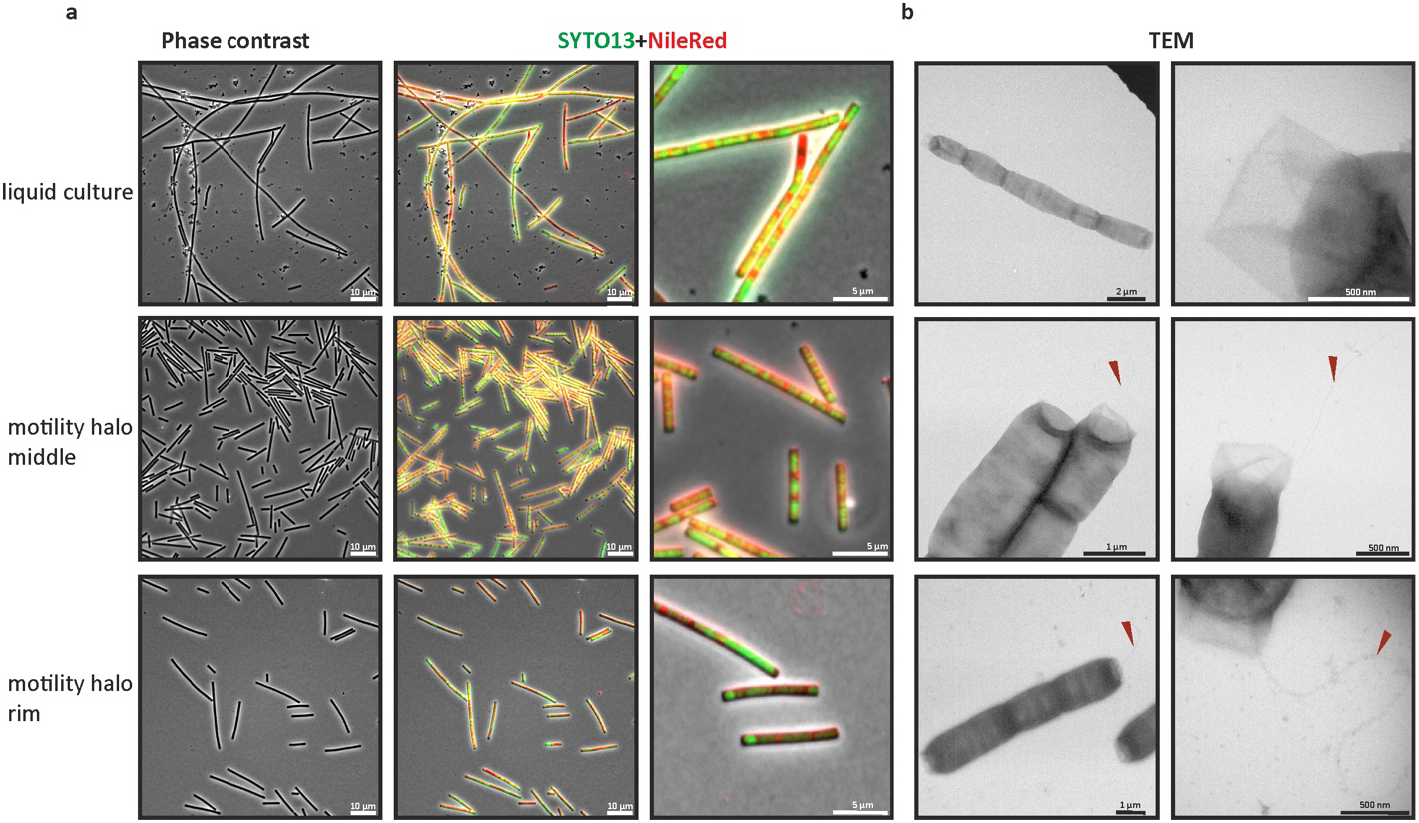
Light, fluorescence, and transmission electron microscopy of the archaellated bacterium *L. aerophila*. A. Light and fluorescence microscopy of *L. aerophila* after labeling with NileRed and SYTO13 from liquid cultures and motility plates. Cell filaments shorten after growth on semi-solid agar plates. The scale bars are 10 µm and 5 µm. B. Transmission electron micrographs of cells. Cells isolated from semi-solid agar plates show distinct polar-located cell surface filaments reminiscent of archaellum filaments indicated by a red arrow. Scale bars are as indicated.

In most Archaea, archaellation is not constitutive ^19,61,63,69,70^, therefore, *L. aerophila* cells were spotted on semi-solid agar plates. After five days, a thin halo appeared and cells from the rim and midpoint of the halo were imaged with light and fluorescence microscopy (Figure 4). Cells from motility plates were mostly shorter (4-10 µm), yet remained multicellular, as shown by fluorescence microscopy (Figure 4). Individual cells of a multicellular filament were separated through a septum (Figure 4a). Electron microscopy imaging of the short filaments of cells revealed surface appendages with 11-12 nm in diameter, exclusively found at the cell poles of shorter multicellular filaments (Figure 4b). Surface appendages were present on cells isolated from the rim of the motility plate as well as from the midpoint (Figure 4b). Additionally, qRT-PCR showed the expression of the archaellin encoding gene *arlB* as well as the archaellum machinery genes in cells from semi-solid agar plates compared to liquid medium grown cells (Supplementary Figure 3).

Given the presence of the archaellum filament, the swimming motility of *L. aerophila* was imaged by time-lapse light microscopy. Swimming cells were analyzed and tracked using TrackMate7^35^. Cells from a motility plate displayed motile behavior with a swimming speed of 10.46 ±6.68 µm/s (Supplementary Video 1). This presents the first actively swimming member of the Chloroflexota phylum and, notably, the first example of a bacterium utilizing an archaellum for motility.

### CryoEM single-particle analysis of the bacterial archaellum filament

To prove that the identified surface structure is assembled by the genes encoded in the archaellum cluster, the filaments were purified. Mass spectrometry indeed validated that ArlB is the main filament protein.

Next, the filaments were subjected to CryoEM single particle analysis. The micrographs contained numerous filaments that were picked using the CryoSPARCs’ implemented filament tracer (Supplementary figure 7). Based on the structural similarity of ArlB from the Euryarchaeum *Methanocaldococcus villosus* to the Alphafold2 predicted structure of ArlB from *L. aerophila*, the helical parameters of the *M. villosus* archaellum were used as initial parameters for helical refinements^28,71^ (108° twist, 5.57 Å rise). These were further refined to 108.14 ° twist and 5.64° rise, resulting in a map with 3.67 Å resolution, which was further improved to 2.7 Å resolution after several rounds of CTF refinement and local motion correction. The final helical parameters of 108.14° twist and 5.64 Å rise were corroborated by helical refinement in CryoSPARC without implying helical parameters and CryoSPARCs own symmetry search job (Supplementary Figure 7,8, Supplementary table 1). The final map revealed a helical structure of the filament with an alpha-helical core decorated with globular domains. Using ModelAngelo, the sequence of ArlB from *L. aerophila* was fitted unambiguously^27,72^. The first 28 amino acids were lacking, and the starting amino acid was isoleucin at position 29. Archaellins possess a signal peptide, which is processed by the class III signal peptidase^8,9^.

*L. aerophila* ArlB has an unusually longer N-terminus compared to archaeal archaellins, but this could be due to wrongly predicting start of the gene. Indeed, ArlB has a class III signal peptide cleavage site, and SignalP6^73^ confidently predicted a cleavage site with shorter N-termini starting at M14 (Supplementary Figure 2). Each ArlB subunit consists of an N-terminal alpha helical tail and a C-terminal globular domain with a characteristic archaellin fold (Figure 5, Supplementary Figure 4). An MSA of bacterial ArlBs showed that the peptidase cleavage site with a consensus cleavage site (G/ITALE) and the hydrophobic part of the signal peptide were highly conserved (Supplementary Figure 1). Similarly to known archaeal archaellum structures, the N-termini of each subunit contributed to the core of the filament interacting with the adjacent subunits through hydrophobic interactions (Figure 5, Supplementary Figure 4).

**Figure 5:**
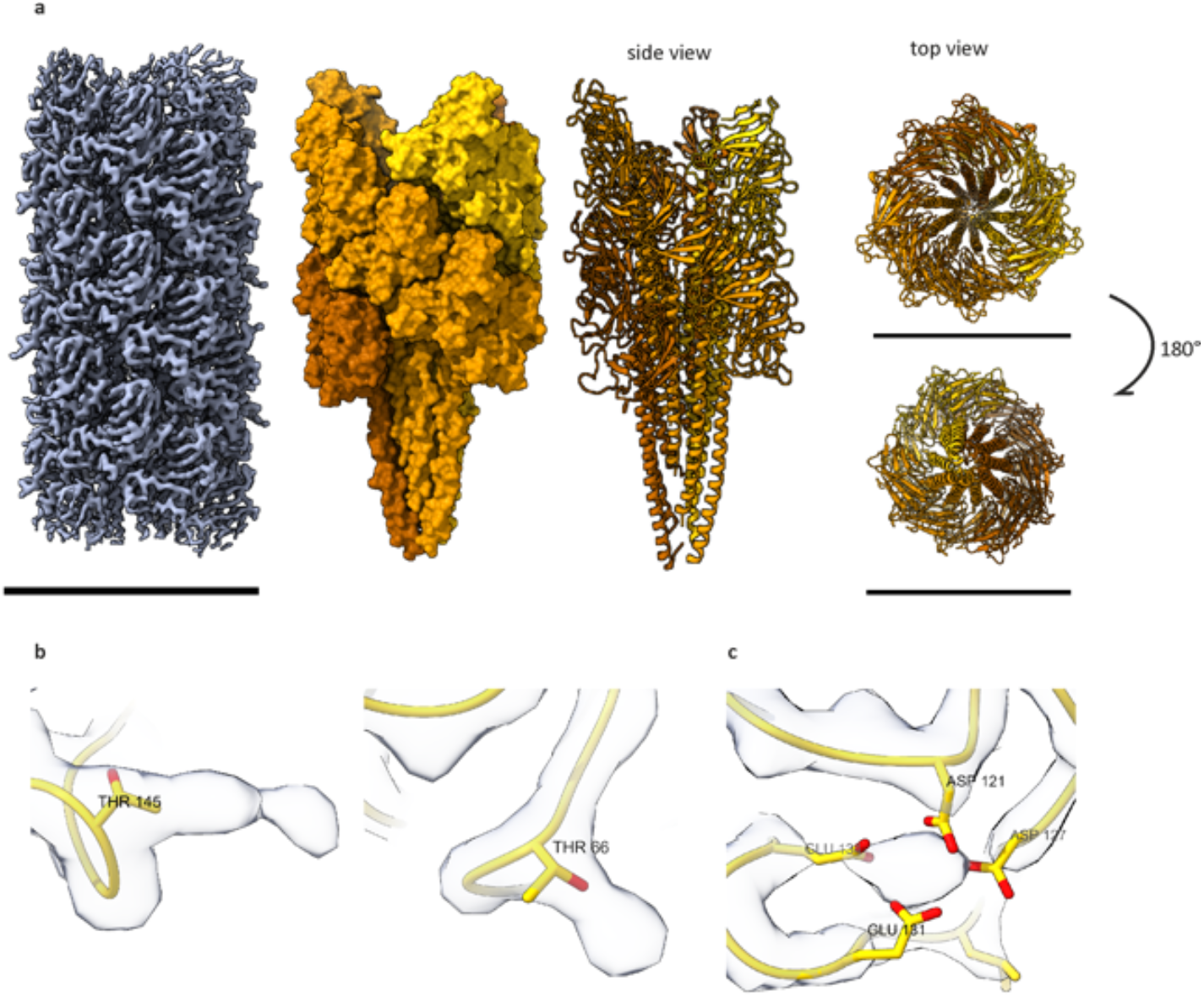
CryoEM structure of the *L*.*aerophila* archaellum. A. cryoEM-derived map at 2.7 Å resolution in surface view with fitted chains in surface and cartoon representation. Cartoon representation of the archaellum structure in side view, top and bottom view showing the core of alpha helices and globular domain facing outwards. Three left-handed strands are colored accordingly. The scale bar is 100 Å. B. Additional dead-end protrusions corresponded to the residues Thr145 and Thr66, likely to be O-glycosylations. C. Each subunit harbors a metal binding site coordinated by the residues D121, D127, E131, E134.

### The bacterial archaellum is remarkably similar to true archaeal archaellum filaments

The architecture of the bacterial archaellum filament was compared with the solved structures of archaeal archaellum filaments from *P. furiosus, Methanospirillum hungatii, Methanococcus voltae* and *Saccharolobus islandicus REY15A*^11,28,74–76^. Despite differences in the helical properties, the bacterial archaellum shows remarkable similarity to its archaeal counterpart (Supplementary Figure 5). All filaments have three strands that are winding upon an imaginary axis in a left-handed manner, thereby defining a three-start helix (Figure 5, Supplementary Figure 5). Each subunit of the filament contributes to the inherent helicity and structure of the filament through tight hydrophobic interactions of the N-terminal alpha helix. The C-terminal globular domain is formed by a beta-sandwich fold following the blueprint of a classical archaellin (Figure 5, Supplementary Figure 5). Structural alignments of the bacterial archaellin to those from archaea revealed a high structural homology to euryarchaeal archaellins lacking the additional C-terminal domain found in *S. islandicus* REY15A ArlB with RMSD under 2 Å (Supplementary Figure 5a,c). Additionally, remarkable conservation was seen in the N-terminal alpha helix of the subunits that determine the filament’s architecture. The globular domain showed higher sequential variability, especially the outward-facing part, a feature that is common to archaeal archaellins (Supplementary Figure 5a,b). Two additional dead-end protrusions were found corresponding to residues T66 and T145. These densities were too large to fit any amino acid residues and might represent additional O-glycosylation of the residues (Figure 5). All archaeal archaellum filaments are highly N-glycosylated, while O-glycosylation is only seen in the archaellum filament of *M. hungatii* and *S. acidocaldarius*^76,77^. In Bacteria, O-glycosylation is commonly found in cell surface filaments such as the pilus in *N. meningitis* or the flagella of *C. jejuni*^78^.

Extra densities were detected in the C-terminal globular domain. The density found cannot be assigned to any amino acid residues and is coordinated by the four residues D121, D127, E134, E131, indicating a probable metal binding site. A multiple sequence alignment of archaellins with the metal binding site revealed the conservation of D121 and E134. Structural alignment of the crystal structure of the *M. jannaschii* archaellin had a highly similar metal coordination (Figure 5, Supplementary Figure 6). ConSurf analysis showed that all four residues are conserved in bacterial ArlBs, indicating conserved metal coordination in the bacterial archaellum similar to euryarchaeal archaellum filaments (Supplementary Figure 6).

**Figure 6:**
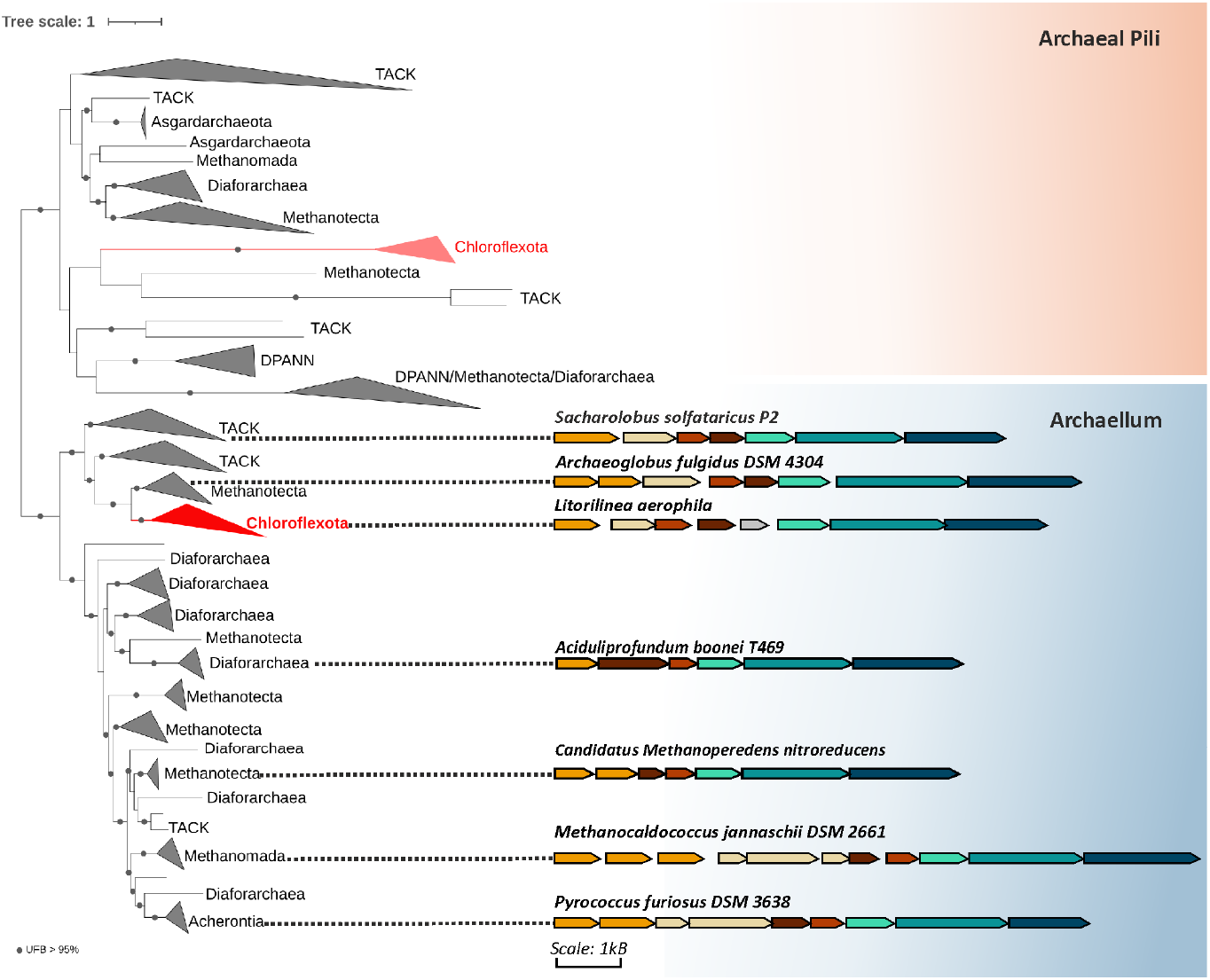
Horizontal gene transfer of the archaellum machinery to Bacteria. Phylogenetic tree of concatenated core archaellum machinery and archaeal pilin subunits. The maximum-likelihood tree was inferred from a supermatrix of 431 sequences and 987 amino acid positions with the model LG+R10+C60. Two horizontal gene transfer events are seen from Archaea to Bacteria. While the first transfer is a remnant archaeal pilus (fainted red), the second transfer led to the diversification of the archaellum machinery in Chloroflexota. Archaellum gene clusters of representative archaellated species are shown colored as in Figure 2 indicating a transfer of an *arl2* cluster to Chloroflexota. Scale bar as indicated.

### Horizontal gene transfer of the archaellum machinery to Chloroflexota

To unravel the origin of the archaellum cluster in Chloroflexota, a concatenation of the core machinery proteins (ArlIJ) present in one copy in the cluster was used to infer a phylogenetic tree of archaeal T4P loci in Archaea and Chloroflexota. By mapping the different subtypes of archaeal T4P according to Makarova et al.^4^ on the resulting phylogeny, we identified two well separated clade (UFB= 100%) (Supplementary Figure 10). While the first clade predominantly contains archaeal pili comprising subtypes as UV pilus, adhesion pilus, bindosome and Epd pilus, the second clade comprises solely archaellum clusters (Figure 6, Supplementary Figure 10). Chloroflexota members are well nested within both archaeal clades, indicating two separate horizontal gene transfer events. The first clade contains two members of Chloroflexota (Anaerolineales order). However, as they lack a pilin subunit these systems are probably incomplete and likely nonfunctional. The second clade contains 232 archaellum loci belonging to Dehalococcoidia and Anaerolineae classes (Figure 6, Supplementary Figure 11). The completeness of the archaellum system within these genomes is striking and indicates a complete transfer of the archaellum cluster to Bacteria. This transfer probably happened from euryarchaeal members belonging to Methanotecta (UFB= 100%) that code for the *arl2* like archaellum cluster to Chloroflexota (Figure 6, Supplementary figure 10). Gene transfers between the two domains of life are frequent, however highly asymmetric with more frequent gene transfers from Bacteria to Archaea than vice versa^79^. Indeed, previous analysis of TFF System in Bacteria and Archaea revealed a horizontal gene transfer of an ancient archaeal pilus to Bacteria led to the bacterial Tad system^3^. The presence of the archaellum machinery in Chloroflexota depicts the second horizontal gene transfer event of a functional cell surface machinery to Bacteria.

## Discussion and Conclusion

The TFF superfamily diversification in procaryotes is driven by adapting existing macromolecular machineries, creating subtypes for specific functions. In Archaea, an ancient TFF machinery was adapted to allow for rotation of a TFF filament, the archaellum^3,7^. Although Archaea typically possess archaella for motility and Bacteria use flagella for this purpose^1^, Hug et al. firstly reported Archaea-specific archaellum genes in metagenomes of Chloroflexota^23^ and Lim et al., reported the presence of the archaellum locus in cultivated members of the SAR202 clade^22^. The archaellum clusters were not widely distributed in Bacteria but were found to be limited to the phylum Chloroflexota. Notably, when archaellum-related genes were identified, flagellar components were absent (Figure 1ab). Indeed, a recent study identified the so far only flagellated Chloroflexota *Tepidiforma thermophila*^58^. Phylogenetic reconstruction indicated that the presence of flagella may have been ancestral in the Chloroflexota, but the flagellum cluster was lost multiple times during the diversification of the phylum, notably due to genome reduction^58,80^. It is plausible that some Bacteria gained the simpler archaellum motility machinery over the complex bacterial flagellum after its loss, and single locus organization of the archaellum machinery may have facilitated multiple horizontal gene transfer events^81^. This has been shown previously through the emergence of the bacterial Tad pilus upon acquisition of an ancient archaeal pilus^3^. Here, an ancient archaeal pilus reminiscent of the methanogenic Epd pilus was acquired by Bacteria and later diversified to the Tad pilus. With this, further components were incorporated as the secretin to accommodate for the diderm envelope^3^.

Our phylogenomic analysis of the archaellum cluster in Archaea and Bacteria revealed two separate horizontal gene transfer events that led to the co-option of the archaellum machinery in Chloroflexota. The first transfer comprises an incomplete archaeal pilus cluster. As a pilin encoding gene is not found within these clusters, it is likely that a remnant machinery comprising the ATPase and membrane protein were acquired through horizontal gene transfer (HGT). The lack of a dedicated pilin subunit probably renders this machinery nonfunctional. The second main HGT can be found within the archaellum specific clade containing all Chloroflexota with an archaellum locus (Figure 6, Supplementary figure 10). The complete archaellum machinery was horizontally transferred to the ancestor of Dehalococcoidia and Anaerolineae in the Chloroflexota. Bacteria and Archaea living in extreme environments tend to share their genes through horizontal gene transfer^82^. However, Chloroflexota with archaellum clusters are present in diverse environments, such as moderate marine, thermophilic, and anaerobic habitats, indicating that environmental factors partly influenced the acquisition and diversification of motility machinery but were not the major driving force. However, it is evident that Chloroflexota took up several genes of archaeal origin. These are mostly enzymes such as the Archaea-specific Mbh hydrogenase or Malate dehydrogenase. It is striking that next to archaellum genes, Chloroflexota from aquifer sediments code of an archaeal type ATP synthase, and one might envision this as a necessary adaptation to the ATP-consuming archaellar machinery^23^.

The presence of archaellum-related genes in Chloroflexota members is indicative of a conserved mechanism of assembly and function in the cell envelope (Figure 2, 3), and indeed, swimming motility is observed (Supplementary Video 1). In *Sulfolobales*, the interaction of the archaellum machinery with the S-layer is essential for the rotary function. Deleting the S-layer genes *slaAB* in *S. islandicus* rendered cells immotile despite having an assembled archaellum. The presence of SlaA enables motility through the interaction of stator proteins ArlFG with the S-layer^15^. Yet, some Archaea display motility despite lacking an S-layer, such as *Oxyplasma meridanum* (Thermoplasmatales)^70^, and this is also the case of Archaea with complex envelope structures as *Methanospirillum hungateii*^83^, suggesting that the archaellum machinery can easily adapt to various cell envelopes. Similarly, the phylum Chloroflexota harbors a wide diversity of cell surface architectures, including single-membraned members with an S-layer, a structure commonly seen in Archaea^58,84–87^. This simple envelope architecture observed in Chloroflexota may have facilitated HGT and the incorporation of the archaellum machinery, likely limiting its diversification to this phylum. The cultivated and archaellated bacterium *L. aerophila* possesses an outer layer resembling the archaeal S-layer and also encodes peptidoglycan synthesis genes (Figure 3, Supplementary Figure 12). This is consistent with the encoded proteins upstream of the archaellum locus, which show partial homology to PilO and a C-terminal LysM domain that specifically binds to peptidoglycan, along with the presence of a PilN counterpart, which have been shown to be essential for the incorporation of bacterial type IV pili into the peptidoglycan layer in bacteria^88^ (Figure 2, Supplementary Figure 9). qRT-PCR has confirmed gene expression in archaellated *L. aerophila*, indicating that these proteins are crucial for the assembly of archaella in peptidoglycan-containing envelopes (Supplementary Figure 3). It is probable that the integration of the archaellum machinery into peptidoglycan-containing envelopes may have involved the co-option of accessory proteins.

The core machinery of the Chloroflexota archaellum as well as the filament architecture is similar to Archaea, indicating that the mechanism for rotation might be conserved. In contrast to Archaea, the bacterial archaellum is O-glycosylated (Figure 5). Although N-glycosylation is more prevalent in Archaea and is crucial for motility in Archaea. While filament architecture remains unchanged, motility is substantially reduced or abolished if glycans are missing or only a truncated glycans are present due to aggregating archaella filaments^89,90^. Bacteria employ both N- and O-glycosylation, and predominantly O-glycosylation of pilins and flagellins, which might have been co-opted for the bacterial archaellum filament^78,91^.

Strikingly, *L. aerophila* seems to link archaellation to alteration of its cell shape. Chains of cells become shorter in later growth phases and are then archaellated. The multicellular filamentous cyanobacterium *Nostoc punctiforme* forms hormogonia, which are differentiated shorter filaments that show gliding motility through type IV pili and secretion of polysaccharides^92,93^. These observations are in striking concordance with what was observed in the well-studied archaeon *Haloferax volcanii*, which is only motile in early growth stages with a rod-shaped phenotype^61^. In *H. volcanii*, positioning of the motility machinery and chemotaxis system is polar similarly to *L. aerophila* and is determined by the oscillation of the ParA/MinD homolog MinD4^94^. *L. aerophila* does encode a CheY with strong structural homology to archaeal and bacterial CheY as well as two ParA/MinD homologs (arCOG00589) (Supplementary Table 3, Supplementary Figure 8). Archaea acquired a bacterial-like chemotaxis system and incorporated an Archaea specific chemotaxis protein CheF to receive the CheY relayed signal at the archaellum motor. Genomic analysis predicted the presence of a complete chemotactic system in Chloroflexota however a CheF homolog is missing^95,96^. Analogously to archaea, Chloroflexota might have evolved another protein component to enable signal transfer from the chemotaxis system to the bacterial archaellum motor complex. Further studies are needed to clarify whether functional chemotaxis system and MinD play a related role in *L. aerophila* and its link to cell shape alteration and positioning of the archaellum machinery.

To conclude, our findings unveil a stunning case of horizontal transfer of the archaellum machinery to Chloroflexota. There, it assembles into a functional rotary filament as observed in Archaea, underscoring the exaptation of what was previously considered to be an archaea-specific machinery, repurposed for identical functional roles.

## Material and methods

If not stated otherwise, all chemicals were either purchased from Roth or Sigma.

### Strains, media, transformation and growth conditions

*L. aerophila* ATCC BAA-2444, DSM25763 (DSMZ) was grown in Difco ™marine broth 2216 medium at 55 °C, at 90 rpm shaking with ambient light in 5 ml plastic tubes or on marine broth agar plates solidified with 1,5 % Bacto Agar (BD).

### RNA isolation and quantitative real-time PCR

For RNA isolation, cells from motility plates or 5 ml of liquid cultures grown for 5 days were harvested and briefly washed with PBS. RNA was isolated using TRIzol reagent followed by phenol-chloroform extraction. Residual DNA was removed by DNase treatment, and the removal of DNA was checked with PCR. cDNA synthesis was performed using the ThermoFisher cDNA synthesis kit (Thermo Fisher). Relative qPCR was performed using qPCRBIO SyGreen® Mix (PCRBioSystems) using cDNA as a template. DNase-treated RNA was used as a non-template control. Fold changes were calculated using the Livak method with *rpoB* as a normalizer^25^.

### Motility plates

0.5% Bacto agar (BD) was dissolved in 400 ml of marine broth medium. 5 µL of a one-day old culture of *L. aerophila* was spotted on the plates. Plates were incubated at 55 °C for 5 days in a sealed plastic box.

### Cell surface filament isolation

The cells of the motility plates were harvested and resuspended in 1x PBS with 2% NaCl. Isolation was done as described in Sivabalasarma et al., 2023^26^. Cell surface filaments were sheared using a blender (Russell Hobbs) or a peristaltic pump (Gilson Minipuls). Cell debris was pelleted by centrifugation at 12 000 x g for 25 min. The cells were pelleted from the supernatant by ultracentrifugation at 200 000 x g for 1 h 10 min. The resulting pellet was resuspended in 500 µl 1xPBS with 2% NaCl. This was applied to 4.5 ml of CsCl2 (0.5 g/ml) dissolved in PBS with 2% NaCl for density gradient centrifugation at 250 000 x g for 16h 30 min. A white band in the upper third was recovered and diluted in 8 ml buffer (1x PBS in 2 % NaCl). This fraction was centrifuged at 250 000 x g for 1h. The resulting pellet contained purified cell surface filaments and was resuspended in 100 µl and stored at −20 ° C.

### Negative-stain electron microscopy

5 µL of cells or purified archaella filaments were applied on freshly glow-discharged 300 mesh carbon/formvar coated copper grid (Plano GmbH). This was incubated for 30 seconds, and excess liquid was blotted away. The grid was washed three times with ddH2O and stained with droplets of 2% uranyl acetate. Imaging was done with a Hitachi HT7800 operated at 100 kV, equipped with an EMSIS Xarosa 20-megapixel CMOS camera.

### Cryo-electron microscopy

3.5 µL of isolated archaella filaments were vitrified on freshly glow-discharged Quantifoil R2/2 grids using a Mark IV Vitrobot (Thermo Fisher). The dataset was collected using a Titan Krios equipped with a Falcon4i and a Selectris energy filter (Gatan). The detector was operated in counting mode at a calibrated pixel size of 0.96Å, corresponding to a magnification of 130 000x. 40-fraction movies were recorded using EPU 3.6 (Thermo Scientific) with an exposure time of 1.95 sec and a total electron dose of 40 e-/Å^2^ at a defocus range of −0.5 to −2 µm. 6110 movies were collected and processed in CryoSPARC v4.6 (Supplementary Table 1)^27^. Briefly, movies were motion corrected, and CTF was estimated using patch.motion and patch.ctf within CryoSPARC. Helical segments were picked with the filament tracer job using 150 Å in diameter and a separation distance of 0.4 diameters between segments. The picked particles were extracted with a fourier-cropped box size of 200 px and two rounds of 2D class averaging were done to remove junk particles. Particles from filament-indicating classes were extracted at an initial box size of 512 px and subjected to 2D class averaging. Selected classes were used to determine the helical parameters using CryoSPARC’s helix refine job. Initial helical parameters of *M. villosus* archaellum were assumed^28^ and refined to 108.09 ° twist and 5.63 Å rise. The obtained map was improved with Global CTF and Local CTF refinement as well as reference-based motion correction. After a final round of helical refinement, the parameters were determined as 108.14° twist and 5.64 Å rise. Local resolution and FSC estimation were performed, and resolution was determined to 2.71 Å.

Helical parameters were further corroborated by running helix refine without implying any helical parameters. Using a symmetry search job within CryoSPARC with a range of 107°-109° twist and 4-6 Å rise 4 possible helical parameters were determined. These parameters were used for a subsequent helix refine job. The obtained map was improved with Global CTF and Local CTF refinement, as well as reference-based motion correction with final helical parameters closely matching the initially obtained ones (Supplementary Figure 7,8).

### Model building and validation

Model building was done using the ModelAngelo build command and the fasta sequence of ArlB1 (WP_141610922.1) from *L. aerophila* was used (Jamali et al., 2024). ModelAngelo fitted the sequence in all well-resolved chains of the archaellum filament map. The model was manually curated and adjusted in Coot^29^. The model was further iteratively refined using phenix.realspace.refine and manually correcting for outliers. Phenix validation with phenix.validation, including phenix.molprobity and Phenix. marriage, was run to determine any rotamers and Ramachandran outliers^30,31^. Molecular models and graphs were generated with ChimeraX^32^. Final validation parameters can be found in Supplementary Table 1.

### Light and fluorescent microscopy

Cells isolated from the rim and middle of the motility halos from the motility plate or grown in liquid medium were harvested and resuspended in 500 µL PBS. Cells were washed with PBS and stained with 5 µl NileRed (5 mg/ml in DMSO) (ThermoFisher) and 1 µL SYTO13 (ThermoFisher). The samples were observed on an agarose pad (1% in PBS) using an inverted Zeiss Axio Observer Z1 phase contrast microscope equipped with a Plan Apochromat 100x 1.4 Oil Ph3 M25 objective controlled via Zeiss Blue v.3.3.89. Image analysis was performed using ImageJ.

### Swimming movies and automated cell tracking

For swimming movies, cells form motility plate were inoculated in filtered marine broth medium and incubated for 90 min at 55 °C while shaking. 1 mL of cells were diluted 1 mL prewarmed filtered marine broth medium and 1.5 mL were transferred into round 0.17-mm Bioptechs delta TPG microscopy dish. Imaging was done using an inverted Zeiss Axio Observer Z1 phase contrast microscope equipped with a Plan Apochromat 100x 1.4 Oil Ph3 M25 objective controlled via Zeiss Blue v.3.3.89 preheated at 55 °C. Movies of swimming cells were recorded for 10 minutes in camera streaming mode. Movies were opened in Fiji^33^ with a time interval of 0.051 seconds. The first five frames of a representative movie were used to train the Weka detector^34^ in Fiji^33^ to detect cells and background signals. This trained Weka model was used to automatically track swimming cells in the full movie using TrackMate7^35^. Low quality tracks, and tracks of multiple cells or background were removed by adjusting detection threshold in TrackMate7. Tracks were refined through visual inspection and manual bridging of gaps within tracks.

### Chloroflexota database and reference phylogeny

We retrieved 9868 genomes annotated as Chloroflexota and available in the NCBI as of June 2024. All downloaded assemblies were taxonomically annotated using GTDB-tk 2.1.1^36^and dereplicated using dRep^37^. We used the threshold of 96% for dereplication to keep representatives from all the species^38^. This resulted in 3780 clusters of Chloroflexota with at least 96% similarities. From each cluster, one representative was selected according to a metric computed based on assembly’s quality scores such as completeness, contamination strain heterogeneity and N50^37^. Next, we used PROKKA^39^ to infer proteins from all the assemblies and build a database combining all Chloroflexota proteomes. To construct a reference species phylogeny of Chloroflexota, we first searched for three conserved and universal markers (IF-2, RpoB, and RpoC) in the database. We used HMMSEARCH^40^ and the pfam domains PF04997 and PF04998 for RpoB; PF04563 and PF04562 for RpoC; and PF11987 for IF-2 and extracted the markers from all Chloroflexota assemblies. We discarded the assemblies with only one marker out of the three and concatenated the markers for the remaining assemblies into a supermatrix of 3543 sequences and 2131 amino acid positions. We inferred a preliminary phylogeny using FastTree2^41^ which was used to sample taxon according to patristic distances using TreeCluster^42^. After two rounds of TreeCluster using length clade= 0.1 and max clade= 0.3 in the first and second round respectively, we selected 443 assemblies, representative of all the taxonomic and phylogenetic diversity of Chloroflexota, and a supermatrix with 443 sequences and 2456 amino acid positions was generated. Finally, a maximum likelihood phylogeny was inferred using IQ-TREE (v2.3.4)^43^ and the evolutionary model LG+C60+I+R10.

### Bacterial protein secretion systems and archaeal TFF detection

To search for TFF and flagellar macromolecular systems, we downloaded the 20 models implemented in the package TXSScan (v1.1.3) dedicated to the genomic detection of bacterial secretion systems and related appendages, including the archaeal pili and archaellum^3,44^. Next, we used MacSyFinder2^45^ and scanned the 3780 genomes of Chloroflexota, as well as 3702 genomes of Archaea and 1048 genomes of Bacteria from our locally maintained databases^46,47^ for the presence of each system. Finally, all the hits identified as best solution by MacSyFinder2 were retrieved and analysed.

### Phylogenetic analyses

Sequences corresponding to the most conserved components of the archaellum were extracted from Chloroflexota database and from a taxonomically balanced subsample of the Archaea database, aligned using MAFFT^48^ with the linsi option, trimmed using trimAl^49^ with the option gappyout, and single gene maximum likelihood phylogenies were generated using IQ-Tree2 (v 2.32.4) with the best-fit evolutionary model selected by BIC criteria^50^.To infer the origin of the archaellum in Chloroflexota, we extracted the sequences corresponding to the core machinery ArlIJ from the archaellum loci comprising only one copy of each gene. Single gene alignments were generated using MAFFT (with the linsi option), trimmed using trimAl (with the option gappyout) and concatenated into a supermatrix of 431 sequences and 987 amino acid positions. Finally, a maximum likelihood phylogeny was generated using IQ-Tree2 and the evolutionary model LG+R10+C60. All the trees were annotated using IToL ^51^.

### Structure prediction analysis

Structure prediction was done with Alphafold3server^52^ (alphafoldserver.com) and visualized through ChimeraX^32^. Structural conservation was analysed through ConSurf WebServer^53^. Gene loci were depicted using GeneGraphics^54^. Weblogo of bacterial ArlBs were created using WeblogoServer^55^. Peptidoglycan synthesis pathway was mapped used KEGG-mapper^56,57^.

## Authors contributions

Major contributions to the concept and design of study (SS, MJ, SVA); Acquisition, analysis or interpretation of data (SS, NT, MJ, CLM, SVA, SG); Writing of the manuscript (SS, NT, CLM, MJ, SVA, SG); Provision of resources and supervision (SVA and SG).

## Acknowledgment

We would like to thank Matthias Boll and the Boll lab for MS analysis. We would also like to thank Bertram Daum and Micha Isupov for helpful suggestions regarding helical reconstruction and model building. We acknowledge the EM facility at the Faculty of Biology at the University of Freiburg for access to their microscopes. The TEM (Hitachi HT7800) was funded by the DFG grant (project number 426849454) and is operated by the University of Freiburg, Faculty of Biology, as a partner unit within the Microscopy and Image Analysis Platform (MIAP) and the Life Imaging Center (LIC), Freiburg. We would also like to thank Stefan Steimle at the Freiburg CryoEM facility for help with cryoEM screening and data collection. Electron microscopy data were collected at the Cryo-EM Facility of the University of Freiburg. The Titan Krios G4 cryo-TEM used for imaging was funded by Deutsche Forschungsgemeinschaft (project no. 506518771) and is operated within the Microscopy and Image Analysis Platform (MIAP), University of Freiburg.

## Data availability

The cryoEM map and the atomic model have been deposited in the Protein Data Bank and EMDB under accession numbers PDB-9I5H and EMDB-52629, respectively. Data used produce our results are provided as supporting data can be found here (https://data.mendeley.com/preview/9999vt8h6h?a=244478e5-a7f2-42fc-bcc5-ff2cef852a8d).

