## Supplemental data for "Horizontal gene transfer of the functional archaellum machinery to Bacteria"

for

References ........ 19

### Extended Data

**Extended Data 1: Distribution of TXSS and TFF systems in Bacteria and Archaea.** Excel file contains the identified TXSS systems of 3780 Chloroflexota genomes, 3702 genomes of Archaea and 1048 bacterial genomes of a locally maintained database using MacSyFinder2[1–4] with genes corresponding to identified secretion system and type IV filament superfamily system.

**Extended Data 2: Horizontal gene transfer of the archaellum machinery to Bacteria.** Mapped clades of archaeal TFF based from Makarova et al., 2016[5]. The upper clades are mainly comprised of archaeal pili, while the lower clades contain archaella machineries. One main horizontal gene transfer of the archaellum machinery from Methanotecta to Chloroflexota led to the diversification of the archaellum machinery in Chloroflexota. ArCOGs per subtype of pili are indicated according to Makarova et al., 2016. Scale bar as indicated.

**Extended Data 3: Swimming movie of *L. aerophila* cells.** Swimming cells over 10 min with 0.05 seconds interval were tracked with TrackMate7[6] and analyzed with Fiji[7]. Scale bar is as indicated.

### Supplementary Figures

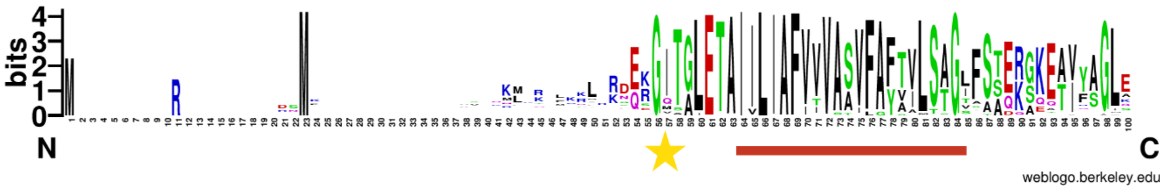

**Supplementary Figure 1: Weblogo of aligned N-terminus of bacterial ArlBs**. The Weblogo depicts the high conservation of the hydrophobic alpha-helical domain and the class III signal peptidase cleavage site. The cleavage site is indicated by a yellow star, and the red bar indicates the hydrophobic domain of the signal peptide.

**a**

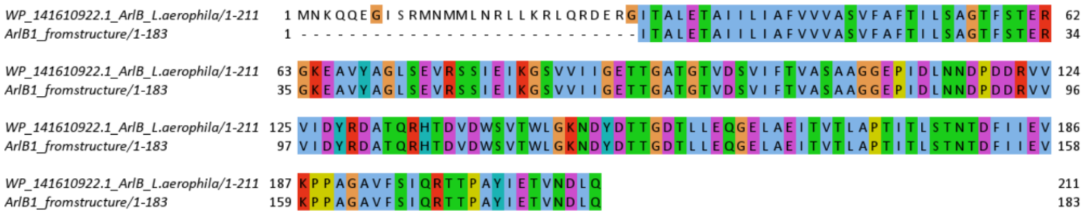

**b**

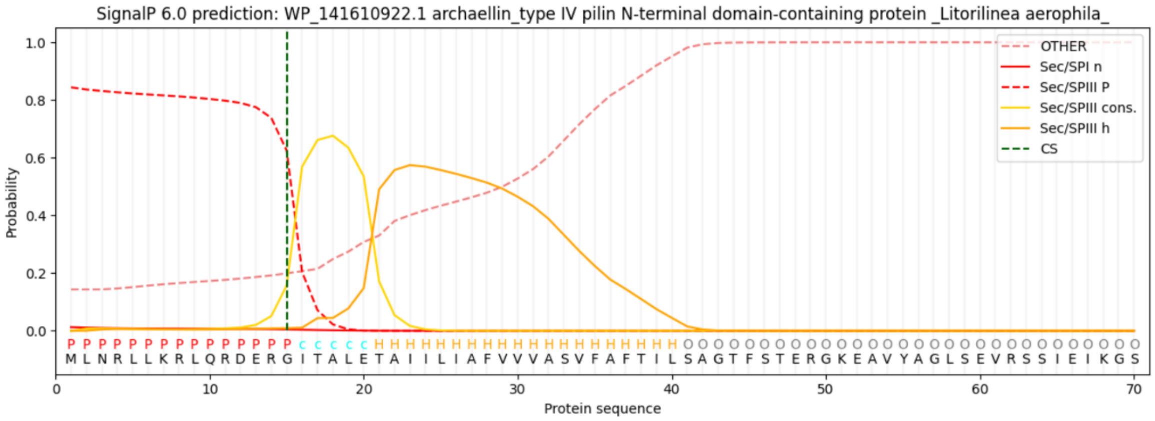

**Supplementary Figure 2: Comparison of structural derived and encoded ArlB** A. The sequence alignment of the experimentally determined ArlB and ArlB derived from the *L. aerophila* genome shows that the ArlB is processed at the predicted class III signal peptidase cleavage motif. B. ArlB SignalP6 prediction with alternative start site at M14 with prediction of a class III signal peptidase cleavage site.

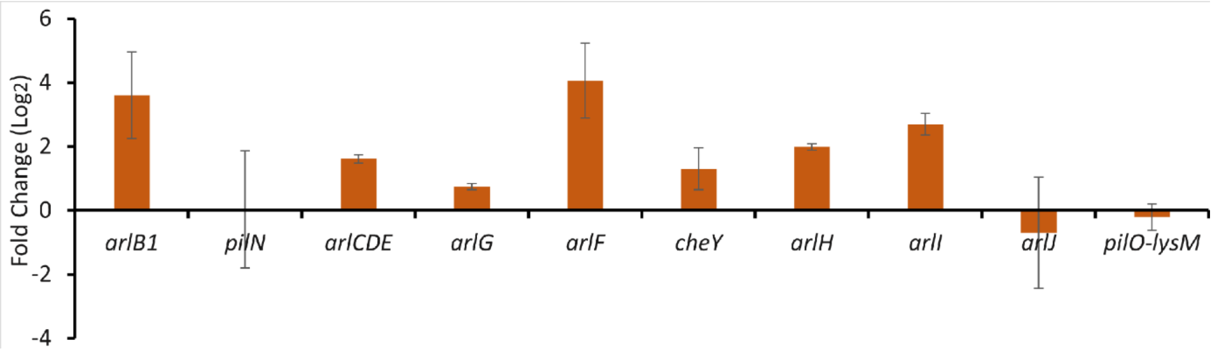

**Supplementary Figure 3: qRT-PCR of archaellum machinery genes.** Relative expression of archaellum machinery genes from *L. aerophila* grown on semi-solid agar plates vs. grown in liquid normalized against rpoB expression.

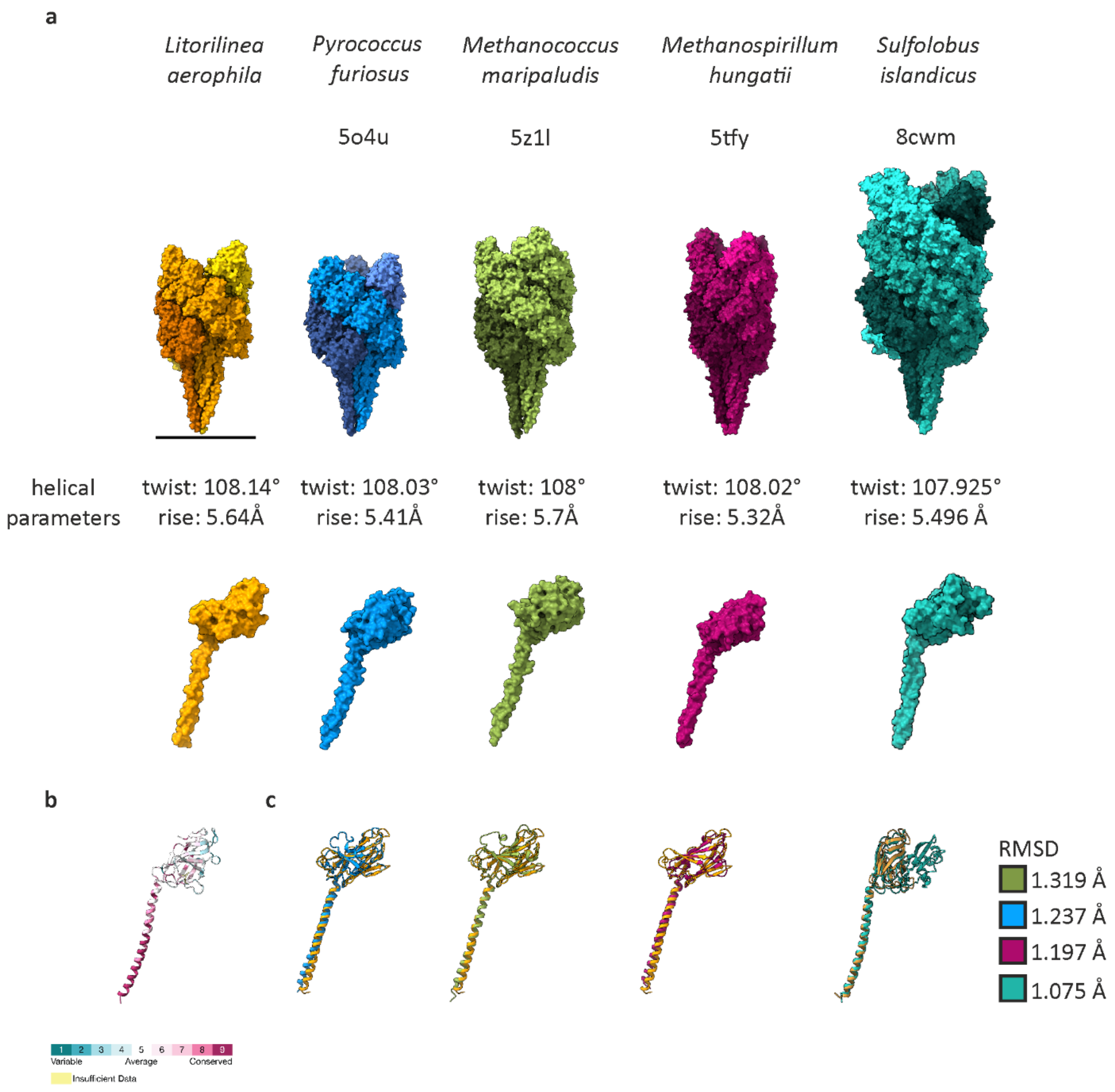

**Supplementary Figure 4: Comparison of the structures of the archaeal archaellum filament with the bacterial archaellum filament**. A. Archaellum filament from *L. aerophila, P. furiosus, M. maripaludis, M. hungatii* and *S. islandicus* in surface representation with indicated helical parameters. Each subunit is shown in a surface representation of the corresponding filament. B. Mapped conservation of bacterial ArlBs on *L. aerophila* ArlB using ConSurf showing a highly conserved N-terminus. C. Structural alignment of *L. aerophila* ArlB1 to archaeal ArlB subunits with indicated RMSD. Scale bar is 100 Å.

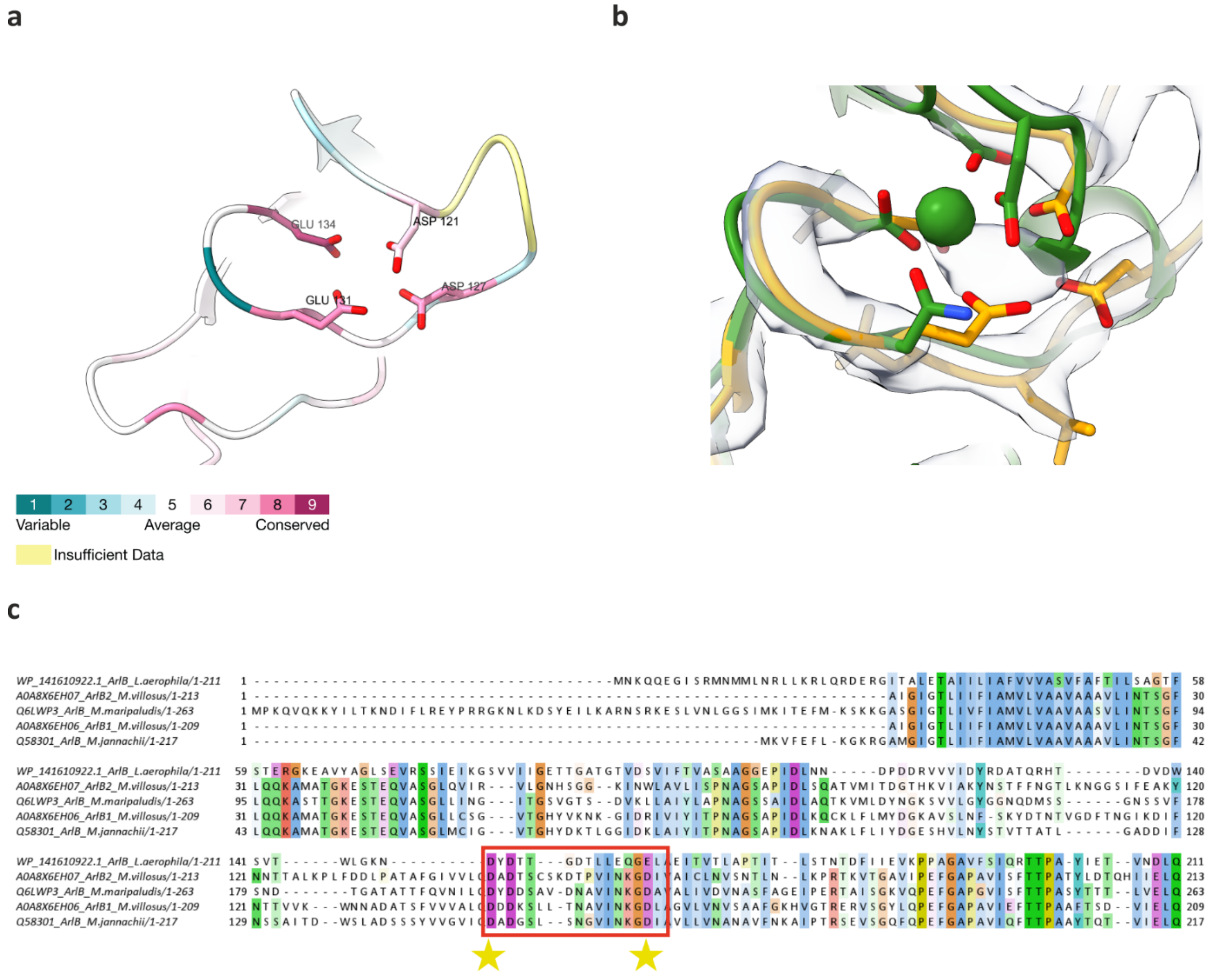

**Supplementary Figure 5: Mapped conservation of residues at the metal binding site using ConSurf.** B. Structural alignment of the metal binding site of archaellin of *M. jannaschii* with coordinated $Ca^{2+}$ ion in green to ArlB of *L. aerophila* in yellow. c. MSA of ArlBs of Euryarchaea with the metal binding site. The metal binding site is indicated in a red box, and two conserved residues coordinating the metal ion are indicated with a star. MSA is colored by conservation.

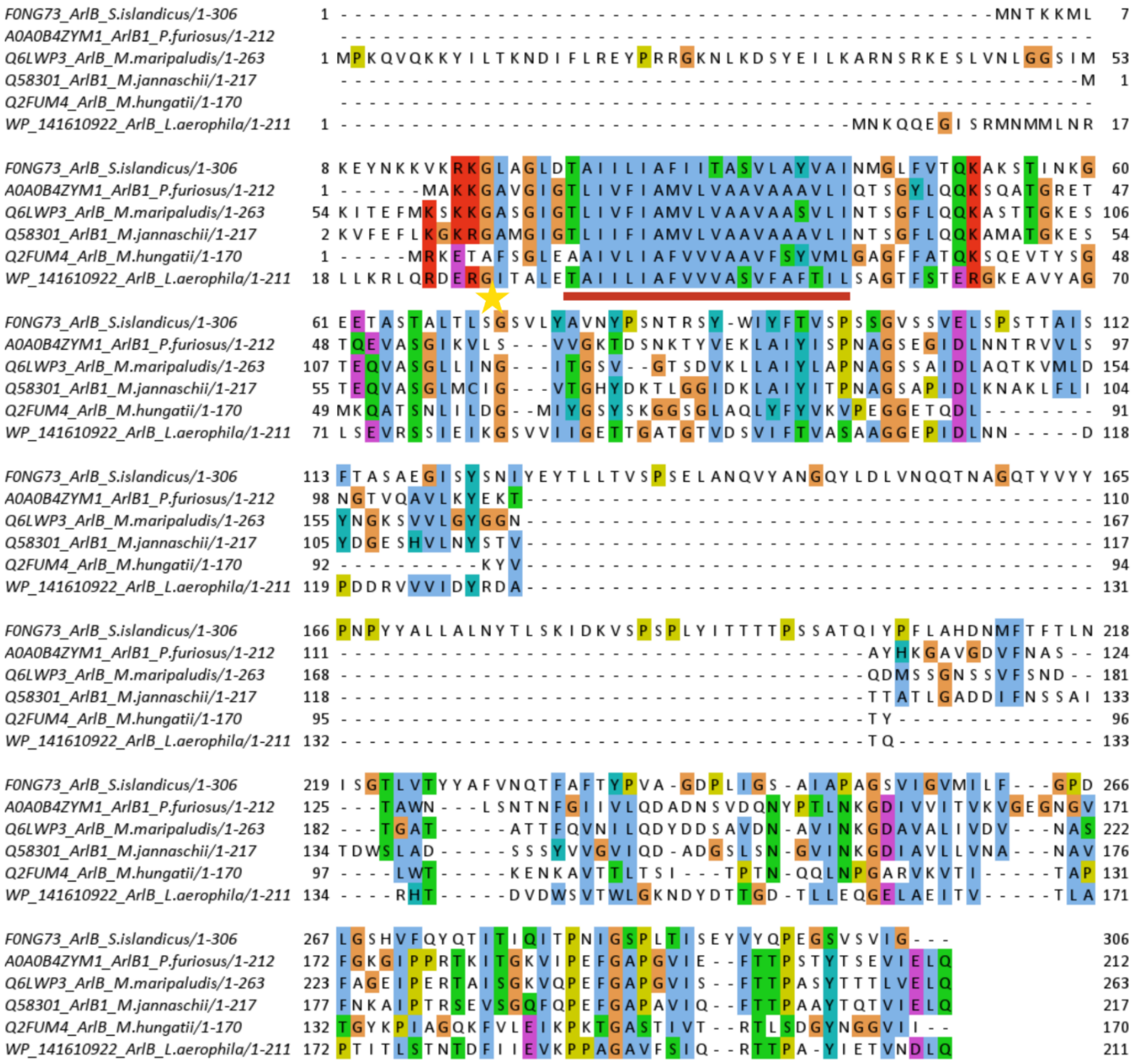

**Supplementary Figure 6: MSA of ArlBs from solved archaellum structures.** The class IIII signal peptide cleavage site is indicated by a yellow star, and the red bar indicates the hydrophobic domain.

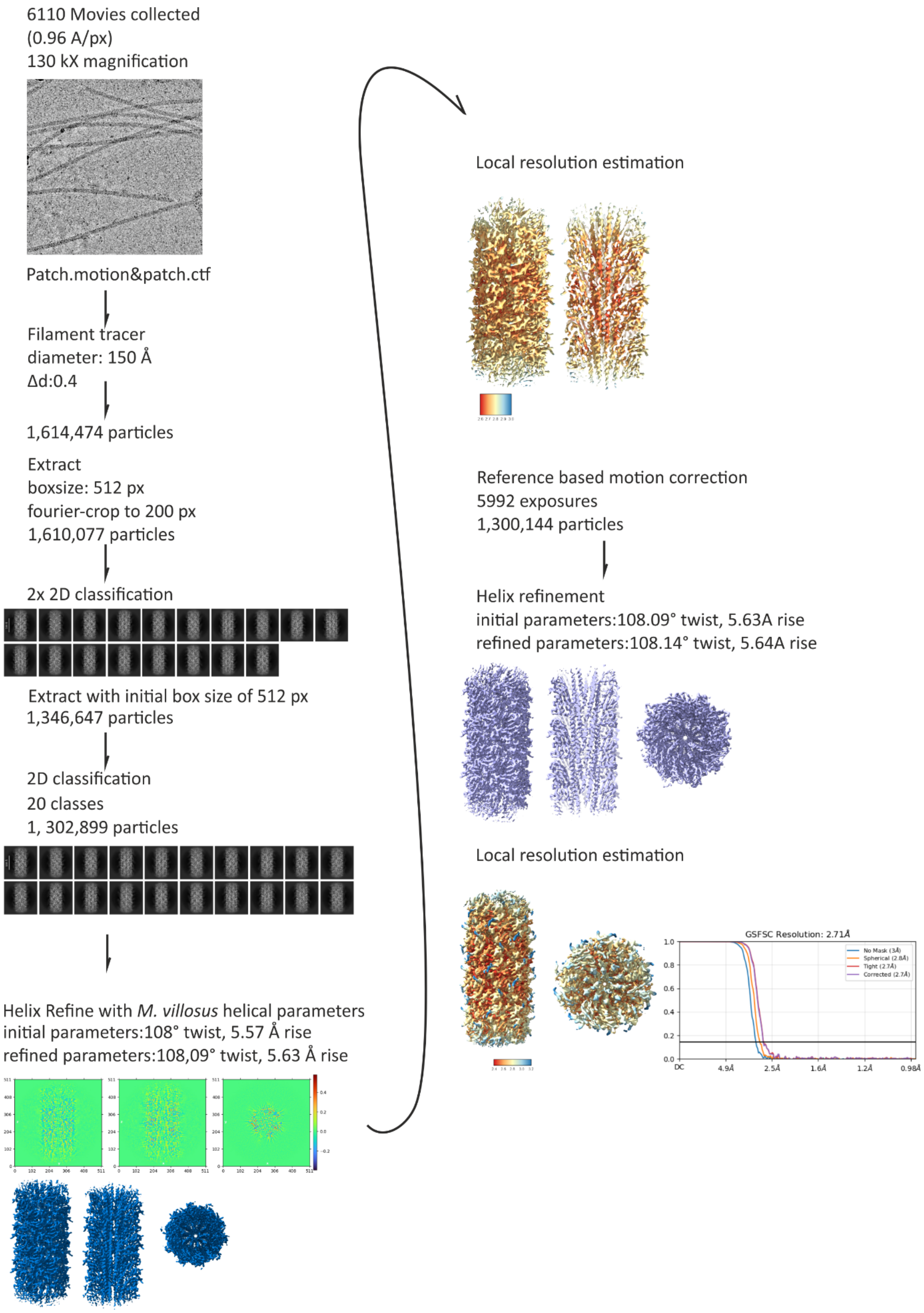
6110 Movies collected
(0.96 A/px)
130 kX magnification
Patch.motion&patch.ctf
Filament tracer
diameter: 150 Å
Δd:0.4
1,614,474 particles
Extract
boxsize: 512 px
fourier-crop to 200 px
1,610,077 particles
2x 2D classification
Extract with initial box size of 512 px
1,346,647 particles
2D classification
20 classes
1, 302,899 particles
Helix Refine with *M. villosus* helical parameters
initial parameters:108° twist, 5.57 Å rise
refined parameters:108,09° twist, 5.63 Å rise
Local resolution estimation
Reference based motion correction
5992 exposures
1,300,144 particles
Helix refinement
initial parameters:108.09° twist, 5.63A rise
refined parameters:108.14° twist, 5.64A rise
Local resolution estimation
GSFSC Resolution: 2.71Å
No Mask (3Å)
Spherical (2.8Å)
Tight (2.7Å)
Corrected (2.7Å)
DC
4.9Å
2.5Å
1.6Å
1.2Å
0.98Å

**Supplementary Figure 7: CryoEM processing workflow with implied helical parameters**.

Helix refine without implied parameters after reference based motion correction

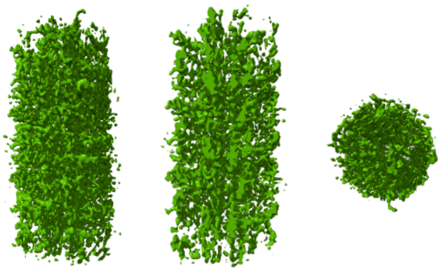

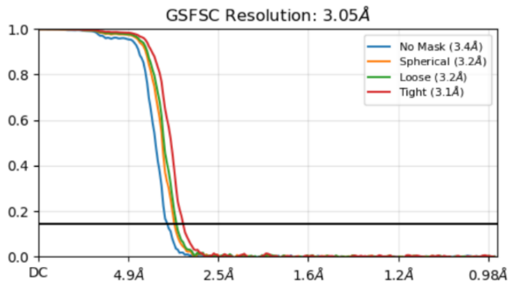

symmetry search job
rise between 4-6 Å
twist: 107°-109°

3 possible parameters

initial parameters:
rise 5.663Å, twist:107.9°

rise: 6Å, twist: 107.9°

rise: 6Å, twist: 107°

refined parameters:
rise 5.62Å, twist108.04°

rise: 5.62Å, twist:107.9°

rise: 5.62Å, twist: 107.94°

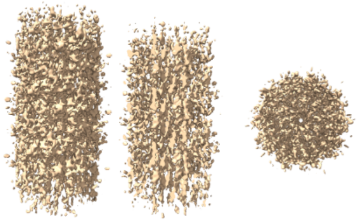

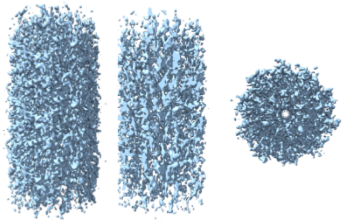

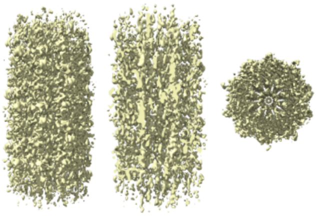

Global and Local CTF refinement

helical refinement

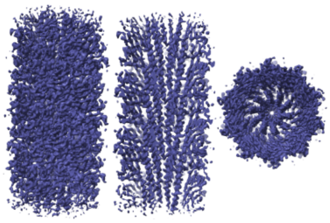

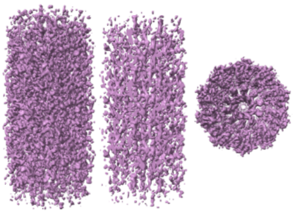

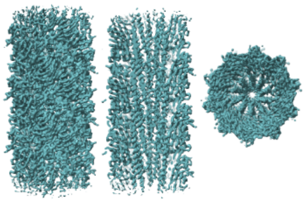

rise: 5.63Å, twist: 108.9°

rise: 5.64Å, twist 107.85°

rise: 5.64Å, twist 108.13°

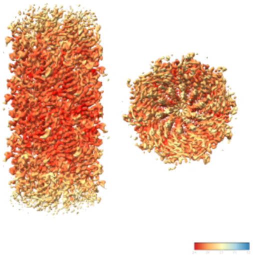

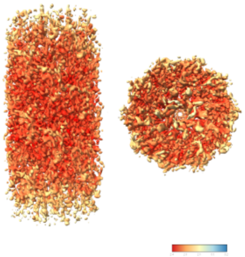

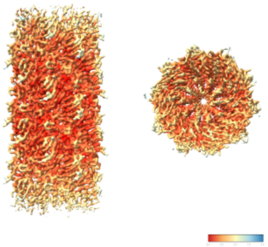

**Supplementary Figure 8: CryoEM processing workflow without implying helical parameters**. A symmetry search job was done after running helix refine without helical parameters and reference-based motion correction.

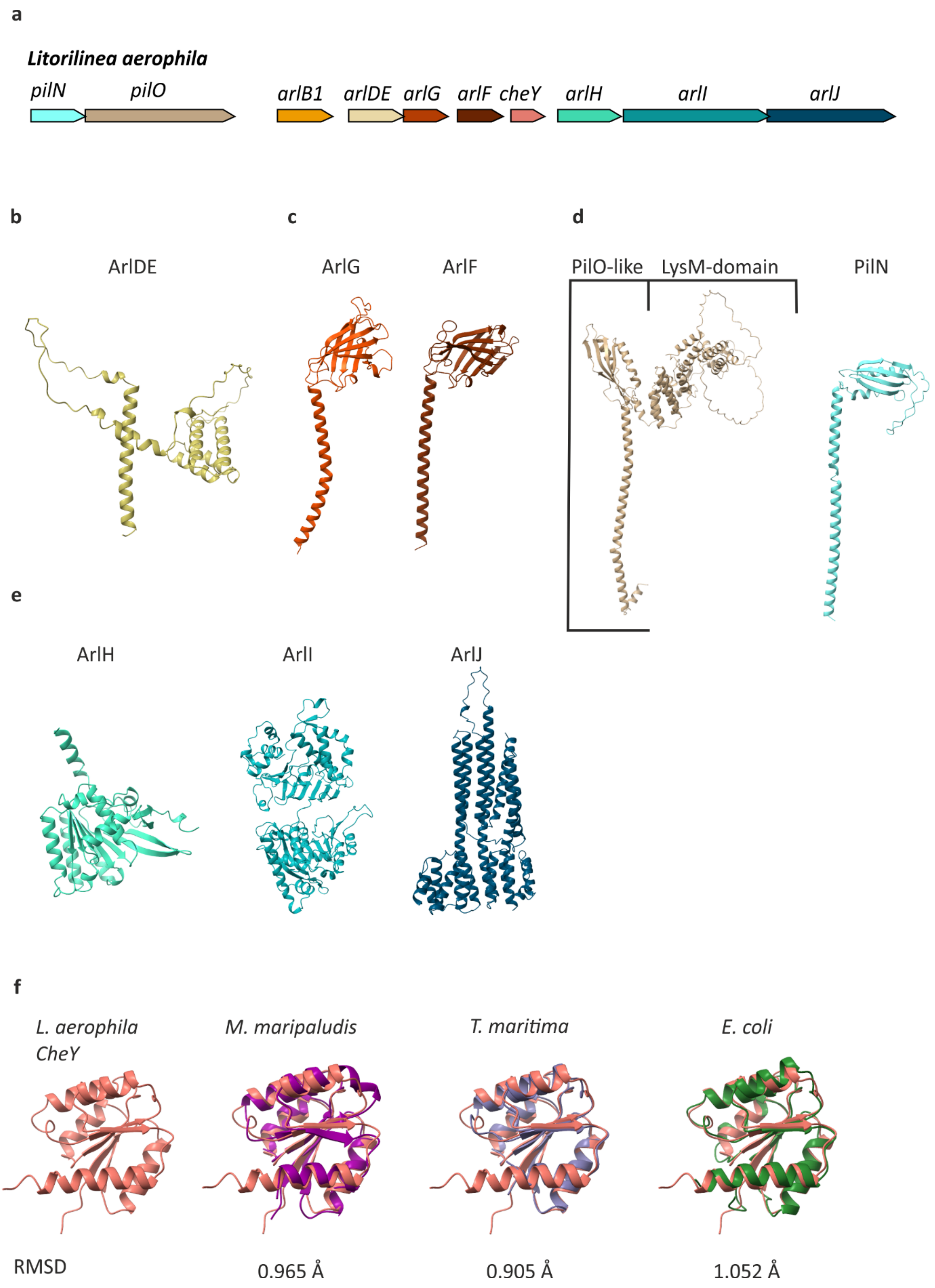

**Supplementary Figure 9: Alphafold3 prediction of the archaellum machinery components of *L. aerophila***. A. shows the genetic locus encoding archaellum-related genes. ArlDE, the archaeal switch complex (b), the stator proteins ArlFG (c), and the core machinery ArlHIJ(e) were predicted and showed similar structures as in motile archaea.

D. L. aerophila encodes for a PilO and PilN homolog. PilO has an additional LysM-domain. F. Structural comparison of L. aerophila CheY found within the archaellum locus and bacterial and archaeal CheY. RMSD is as indicated. The colors of structures correspond to the colors in A.

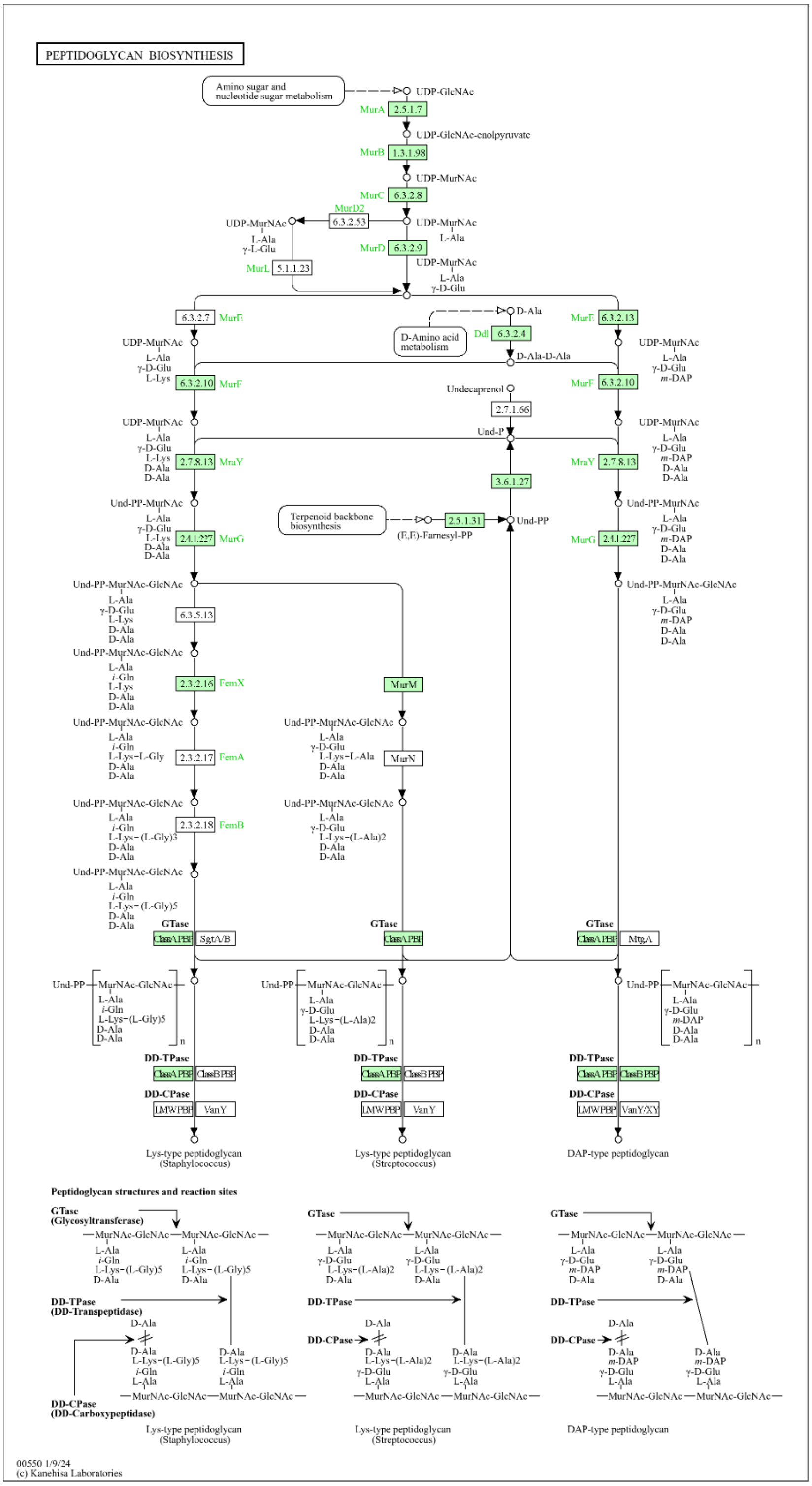

**Supplementary Figure 10: Supplementary Figure 12: Mapped peptidoglycan synthesis pathway in  using KEGG-mapper.**

### Supplementary tables

#### Table 1: CryoEM data collection and processing

**Cryo-EM data collection, refinement and validation statistics**

| | **Bacterial archaellum filament from *Litorilinea aerophila* (EMDB-**52629**)**<br><br>**(PDB** 9I5H**)** |
|---|---|
| **Data collection and processing** | |
| **Magnification** | 130k |
| **Voltage (kV)** | 300 |
| **Electron exposure (e-/Å²)** | 40 |
| **Defocus range (µm)** | -0.5—2.0 µm |
| **Pixel size (Å)** | 0.96 A |
| **Symmetry imposed** | Twist: 108°, Rise: 5.57Å |
| **Symmetry final** | Twist: 108.14°, Rise: 5.64Å |
| **Initial particle images (no.)** | 6110 |
| **Final particle images (no.)** | 6110 |
| **Map resolution (Å)** | 2.71A |
| **FSC threshold** | 0.143 |
| **Map resolution range (Å)** | 2.6-3.0 |
| **Refinement** | |
| **Initial model used** | Ab initio |
| **Model refinement resolution (Å)** | 2.7 |
| **Model composition** | 23316 atoms |
| **Protein residues** | 3110 |
| **Bonds (R.M.S. deviations)** | |
| **Length (Å)** | 0.008 |
| **Angles (Å)** | 0.671 |
| **MolProbity Score** | 1.67 |
| **Clash Score** | 6.03 |
| **Ramachandran plot (%)** | |
| **Outliers** | 0.03 |
| **Allowed** | 4.78 |
| **Favored** | 95.19 |
| **CaBLAM outliers (%)** | 1.64 |
| **C-beta outliers (%)** | NA |
| **CC/masked** | 0.88 |

**Table 2: Primers used for qRT-PCR**

| Primer | Sequence 5´-3´ | Purpose |
|---|---|---|
| 13385 | CATAGTGGCATATTCGGAGACC | qRT-PCR primer fw for *L. aerophila* FKZ61_RS14900-*arlB2* |
| 13386 | ATCTCCTTCACCTCCCTCAA | qRT-PCR primer rev for *L. aerophila* FKZ61_RS14900-*arlB2* |
| 13387 | GTGTCACCTGTGGTATCGTAAT | qRT-PCR primer fw for *L. aerophila* FKZ61_RS14890-*arlB1* |
| 13388 | TCTGAACAATGACCCGGATG | qRT-PCR primer rev for *L. aerophila* FKZ61_RS14890_*arlB1* |
| 14305 | CGTTGCCATTGGTGGATTTG | qRT-PCR primer fw for *L. aerophila* FKZ61_RS14885-*arlCDE* |
| 14306 | GGGAATTTGGCCTGACAGAA | qRT-PCR primer rev for *L. aerophila* FKZ61_RS14885-*arlCDE* |
| 14307 | TCCTTGTAAGGAATGCGGATAAA | qRT_PCR primer fw for *L. aerophila* FKZ_RS14880-*arlG* |
| 14308 | CGACATCTTCGTATGGGTCAAA | qRT-PCR primer rev for *L. aerophila* FKZ_RS14880-*arlG* |
| 14309 | CCGTTAATTGATTGGTGGTGTT | qRT_PCR primer fw for *L. aerophila* FKZ_RS14875-*arlF* |
| 14310 | GTGTATGAGCCGGACATCTT | qRT-PCR primer rev for *L. aerophila* FKZ_RS14875-arl*F* |
| 14311 | GTCCATGCCAGGTAGAACAAT | qRT_PCR primer fw for *L. aerophila* FKZ_RS14870-*cheY* |
| 14312 | GGAAGGCTATCAAGTCCACAC | qRT-PCR primer rev for *L. aerophila* FKZ_RS14870-*cheY* |
| 14313 | GCGCCTTACTGATGGGAATAA | qRT-PCR primer fw for *L. aerophila* FKZ_RS14865-*arlH* |
| 14314 | CTGATCAAGGCCATGGAAGT | qRT-PCR primer rev for *L. aerophila* FKZ_RS14865-*arlH* |
| 14315 | GCGGATCGATGGGAATGTAATA | qRT-PCR primer fw for *L. aerophila* FKZ_RS14860-*arlI* |
| 14316 | GGGTTGAAGGTCCGTAATCTC | qRT-PCR primer rev for *L. aerophila* FKZ_RS14860-*arlI* |
| 14317 | CCCATCACCCAGAGGATAAATG | qRT-PCR primer fw for *L. aerophila* FKZ_RS14855-*arlJ* |
| 14318 | GAAGGAATACGAACGGGATCTG | qRT-PCR primer rev for *L. aerophila* FKZ_RS14855-*arlJ* |
| 14321 | CGTCGCTGTTGCTCTCTT | qRT-PCR primer fw for *L. aerophila pilO-* |
| 14322 | CGGCCATTCAGGAACTCTT | qRT-PCR primer rev for *L. aerophila pilO* |
| 14303 | ACGTCCACTACAGCCATTAC | qRT-PCR primer fw for *L. aerophila* FKZ_RS10530-*rpoB* |
| 14304 | GCGTCTCAATGAAGCCAAAG | qRT-PCR primer rev for *L. aerophila* FKZ_RS10530-*rpoB* |
