## Supplementary figures and images for "Horizontal gene transfer of the functional archaellum machinery to Bacteria"

### Extended Data 2

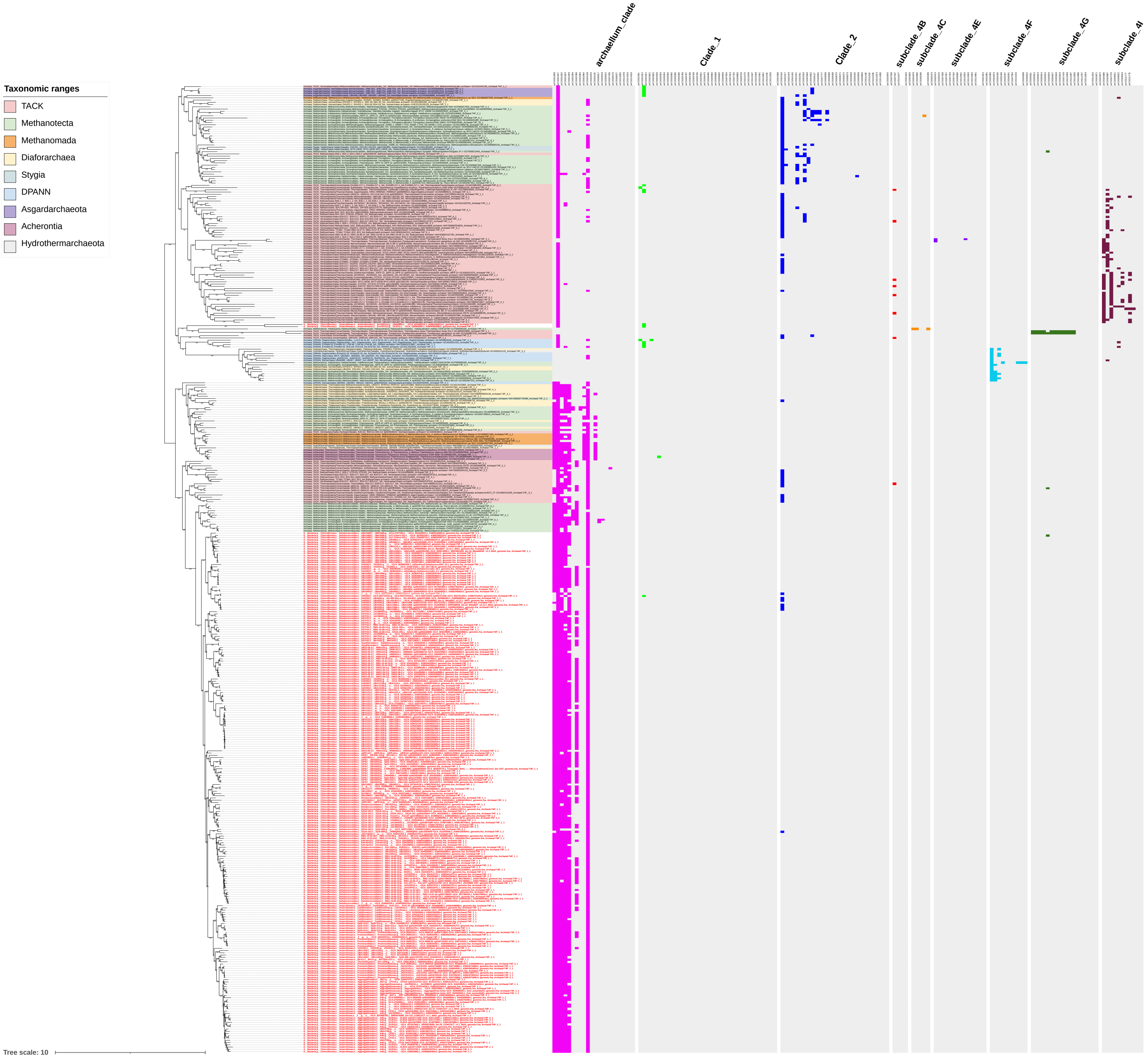
